# Gene Function Revealed at the Moment of Stochastic Gene Silencing

**DOI:** 10.1101/2024.07.16.603770

**Authors:** Shreyan Gupta, James J. Cai

**Author notes:** Corresponding author: James J. Cai, Department of Veterinary Integrative Biosciences, Texas A&M University, College Station, TX 77843, USA.

## Abstract

Gene expression is a dynamic and stochastic process characterized by transcriptional bursting followed by periods of silence. Single-cell RNA sequencing (scRNA-seq) is a powerful tool to measure transcriptional bursting and silencing at the individual cell level. In this study, we introduce the single-cell Stochastic Gene Silencing (scSGS) method, which leverages the natural variability in single-cell gene expression to decipher gene function. For a target gene *g* under investigation, scSGS classifies cells into transcriptionally active (*g*+) and silenced (*g*-) samples. It then compares these cell samples to identify differentially expressed genes, referred to as SGS-responsive genes, which are used to infer the function of the target gene *g*. Analysis of real data demonstrates that scSGS can reveal regulatory relationships up- and downstream of target genes, circumventing the survivorship bias that often affects gene knockout and perturbation studies. scSGS thus offers an efficient approach for gene function prediction, with significant potential to reduce the use of genetically modified animals in gene function research.

## Introduction

During WWII, researchers analyzed bullet damage on returning airplanes to decide where to reinforce armor. But statistician Abraham Wald noted a flaw: they only studied surviving planes, overlooking those shot down. The bullet holes did not mark weak spots, but areas where planes could withstand hits. This was survivorship bias, skewing conclusions by focusing solely on survivors. This idea of survivorship bias is also relevant in the field of genetics and molecular biology. In clinical biomarker studies, for example, geriatric and frail patients are often excluded from studies due to surgical limitations, despite being the ones most likely to have more aggressive forms of glioblastoma (see review ^1^). Ironically, these patients would have been ideal candidates for research. By excluding them, the data collected was not representative of the entire population, introducing limitations for downstream analysis. This type of selection bias is not unique to clinical studies. In genetic research, a similar issue arises when studying the function of a particular gene. Researchers often knock out (KO) the gene of interest and compare the KO sample to a wild-type (WT) sample. In this process, they essentially examine the “survivors”—the cells that remain functional despite the knockout. The cells that die due to the genetic perturbation are disregarded, much like how certain patient groups are excluded from clinical studies. In both cases, the data analyzed does not fully represent the entire system, whether it’s the patient population or the genetic network. Just as Wald cautioned against focusing only on the surviving planes, it would be inadvisable to draw conclusions from samples in which the function of the target gene remains. The bias can skew the interpretation of results, leading to an incomplete understanding of the impact of the genetic perturbation.

Stochasticity in gene expression provides a way to bypass the limitations of survivorship bias that plague gene knockout and perturbation studies. Gene expression in cells is a dynamic phenomenon, modulated by extra-cellular and intra-cellular environmental factors, leading to noise and stochasticity in the data ^2-6^. One such factor leading to stochasticity in gene expression is transcriptional bursting ^7,8^. This term refers to the phenomenon where genes are transcribed in short bursts followed by periods of inactivity or silence, leading to fluctuations in mRNA levels. This variability in gene expression can arise from various sources such as stochasticity in transcription factor binding, biological rhythms, chromatin accessibility, and RNA polymerase activity ^9,10^. The transcriptional machinery of inactive (silent) cells differs from that of cells exhibiting active bursting for a particular target gene. Single-cell RNA sequencing (scRNA-seq) has revolutionized the field of transcriptomics, enabling us to quantify the RNA-expression profiles of individual cells. By profiling the transcriptomes of individual cells, scRNA-seq can reveal the heterogeneity in gene expression dynamics, allowing researchers to study the underlying mechanisms and regulatory networks driving transcriptional bursting.

Dropouts in scRNA-seq data are instances where mRNA molecules from a gene are not captured during the sequencing process, most likely due to low expression levels, resulting in zero values for that gene in that cell. Even though dropouts in scRNA-seq data are a huge source of technical noise, several studies have shown functional associations between dropouts and biological function ^11-13^. The term “dropout” in scRNA-seq is often misleading, as it implies missing data when in reality, it reflects a true lack of or low mRNA expression. Dropouts are a direct representation of gene expression levels within cells. Ignoring dropouts by simply attributing them to measurement leaks overlooks important transcriptional dynamics and can lead to a misrepresentation of the underlying biology. We strongly concur with Sarkar and Stephen ^14^ that scRNA-seq analysis should be approached through two distinct models: 1) an expression model that studies the variability of true gene expression across individual cells and 2) the measurement error model which accounts for technical errors arising from the sequencing process. With this framework in mind, our study exclusively employs the expression model to quantify gene expression levels. Given the transient and volatile nature of mRNA molecules, viable cells with zero expression of the target gene are likely in a silenced transcriptional state, while cells expressing the target gene are likely in an active transcriptional state.

We hypothesize that the cells that are silenced for a gene of interest (that transiently stop expressing it) are functionally distinct. These cells can be identified by the dropout pattern of scRNA-seq data. The cells with zero expression of the target gene are also likely to show a change in expression for closely related genes. Even after splitting the cells into active and silent samples, due to the high number of cells captured by scRNA-seq, we have enough statistical power to perform statistical tests that compare the mean expression changes. This stochastic naturally occurring phenomenon can be imagined as a miniature natural genomic perturbation experiment that is free from survivorship bias. Thus, these small changes in gene expression between active and silent cells are significant and can be used for functional analysis of the target gene.

In this study, we present single-cell Stochastic Gene Silencing (scSGS), an scRNA-seq analysis framework for deciphering natural transcriptional bursting patterns of genes and their associated biological functions. scSGS uses only wildtype (WT) scRNA-seq data and classifies the cell sample into transcriptionally active and silenced subsets based on the binarized expression pattern of the target gene. SGS-responsive genes are identified by comparing the active cells with the silenced cells using the non-parametric Wilcoxon rank sum test. The SGS-responsive genes are then used for functional enrichment analysis to predict the biological impact of that gene on the system. scSGS is an unsupervised de novo gene function inference tool and does not require prior knowledge of gene regulation or biological mechanisms.

The rest of the article is structured as follows: We first present an overview of the scSGS analysis framework. Then, we use publicly available real scRNA-seq datasets to test and validate the biological significance of the SGS-responsive genes. Next, to test the robustness and scalability of the framework, we run scSGS on one common cell type from three datasets of varied sizes and sequencing depths. Then, to compare the gene relation inference capabilities of scSGS, we tested it against the widely adopted Pearson’s and Spearman’s correlation metrics for co-expression analysis. Finally, we compare the performance of scSGS with that of an existing *in silico* gene perturbation prediction tool.

## Results

### The scSGS Framework

The scSGS analysis framework is depicted in **Fig. 1**. The analysis was performed on the scRNA-seq gene expression profiles of WT samples of our interest. The pipeline starts with the preprocessing of scRNA-seq data to filter out cells and genes with low expression profiles.

**Fig. 1.**
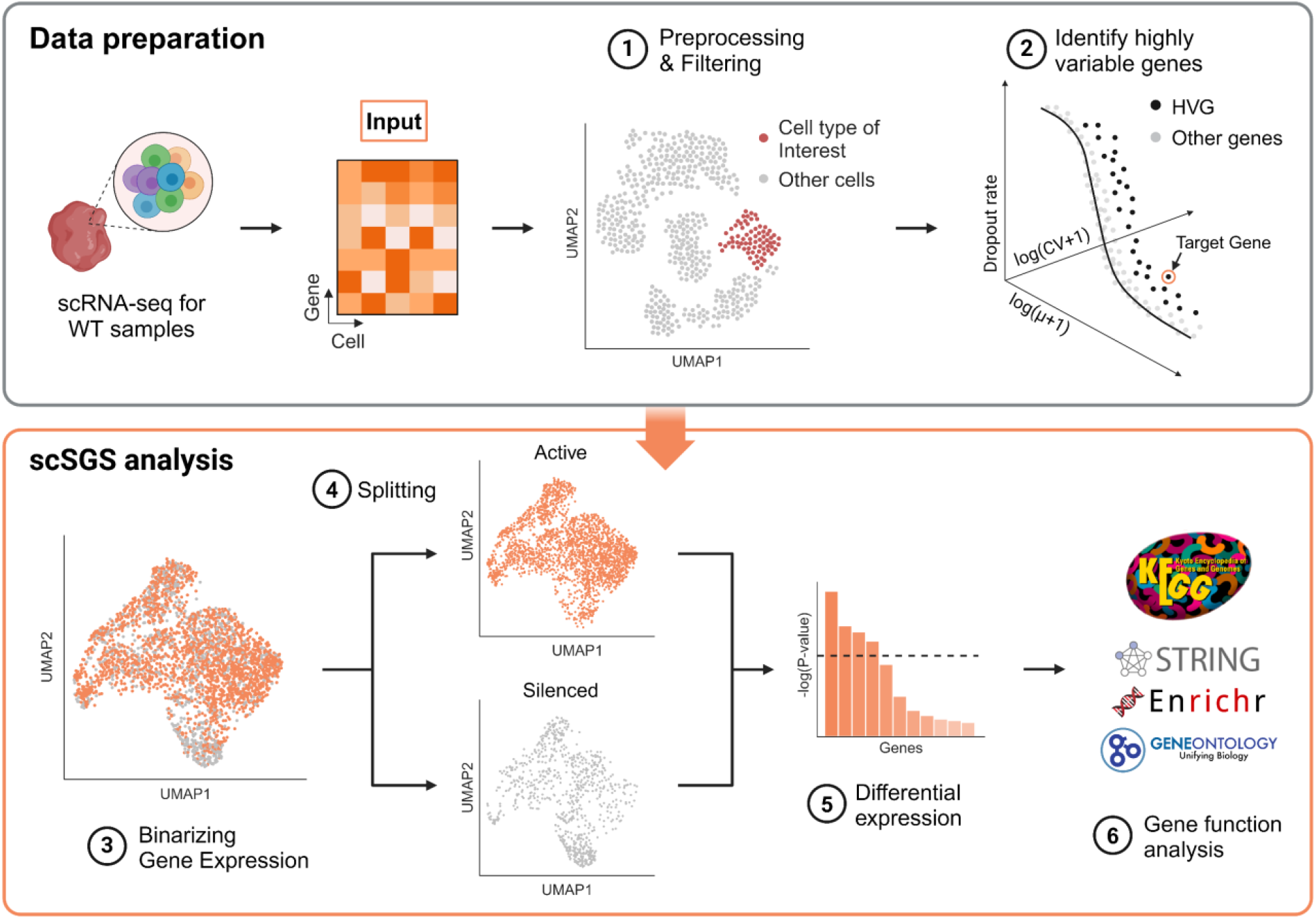
The pipeline contains six steps: (1) pre-processing scRNA-seq data, (2) identifying target genes using Spline-HVGs, (3) binarizing gene expression of the target gene, (4) splitting data based on the expression of the target gene, (5) performing differential expression analysis to compare split subsets, and (6) performing functional annotation and network analysis.

Doing so ensures that the cells left in the matrix are viable and functional. The cell types are annotated using canonical cell type markers from the ScType ^15^ database. Then, cells corresponding to the cell type or state of interest are extracted. Taking the dropout rate and variability in gene expression into consideration, the scSGS pipeline uses a three-dimensional spline-based highly variable gene identification algorithm to identify suitable genes for the analysis. The gene expression of the target gene is then binarized (*G*_*Bin*_) and used as a classifier to split the scRNA-seq count matrix. The cells that express the target gene (*G*_*Bin*_ = 1) are allotted to the active subset, and cells with 0 expression of the target gene (*G*_*Bin*_ = 0) are allotted to the silenced subset. The gene expression profiles from the active and the silenced matrices are normalized and then compared using the non-parametric Wilcoxon rank-sum statistical test. The average log2FoldChange is also calculated to infer the direction and intensity of the SGS response. The lower the P-value of a gene, the greater the association with the target gene. Significantly SGS-responsive genes are identified using a false discovery rate (*FDR)* cutoff of < 0.01. The enriched functions of these significantly SGS-responsive genes are used to predict the biological function of our target gene.

### Real data scSGS analysis unveils *Ccr2* function in glioblastoma

scSGS, as a gene function inference tool, is expected to recapitulate some discoveries from *in vivo* KO experiments and further provide novel gene links. To test the inference power of scSGS, we compared the SGS-responsive genes using only WT samples with *in vivo* KO experiments. We collected our first scRNA-seq dataset from an *in vivo* KO study performed by Pombo Antunes *et al*. ^16^ on the immune landscape of glioblastoma in WT and *Ccr2*^-*/*-^ mice (**Fig. 2a**). The scRNA-seq data was preprocessed and filtered. We preserved the cell type annotations generated by Pombo Antunes *et al*.

**Fig. 2.**
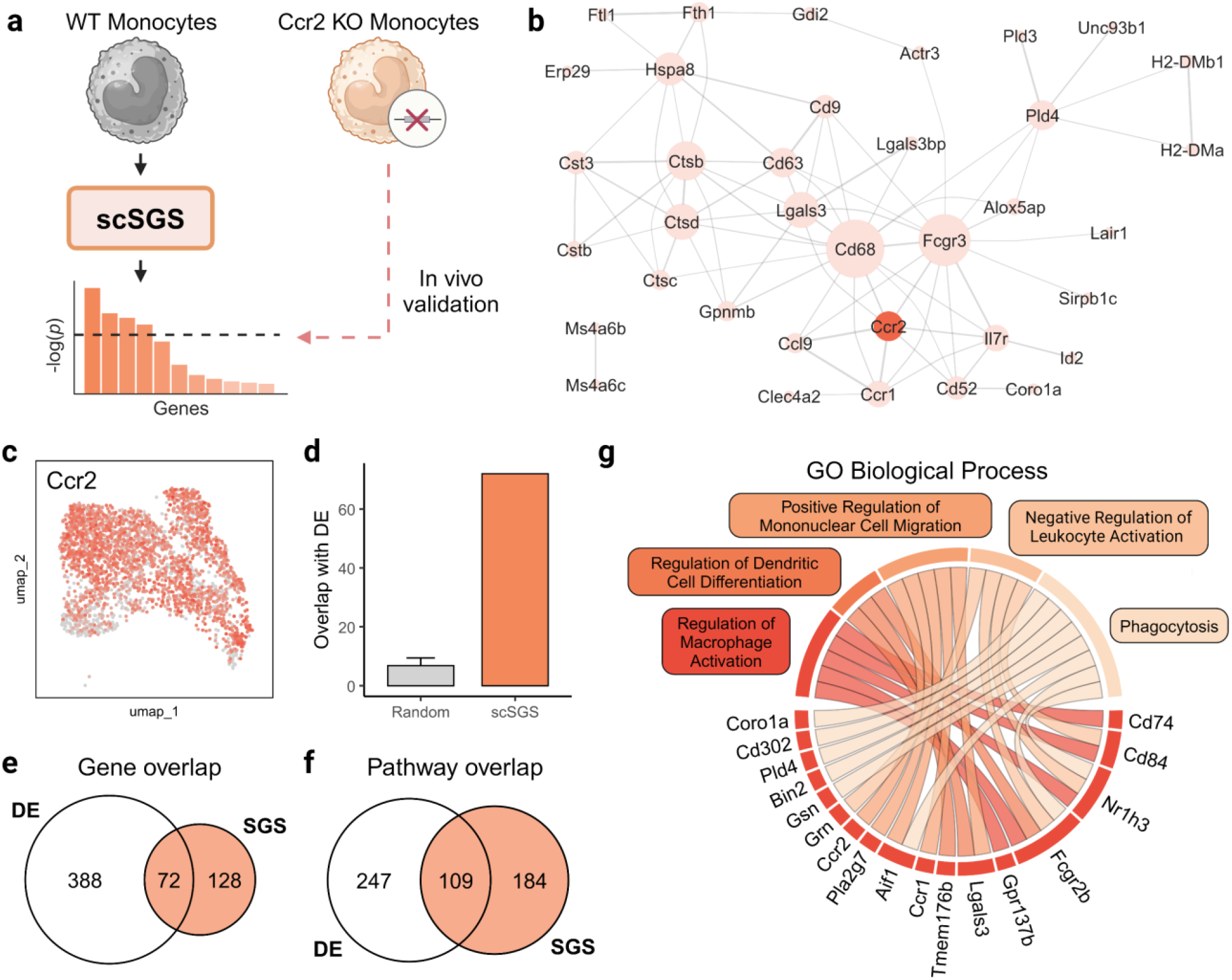
SGS-responsive genes inferred from *Ccr2* perturbation. (a) Experimental design of scSGS analysis on WT and *Ccr2*-KO monocytes from mouse glioblastoma. (b) STRING interaction network of top 50 SGS-responsive genes. Edge thickness indicates the strength of data support and node size indicates node degree. (c) *Ccr2* expression in WT monocytes only. (d) The intersection of 200 randomly selected genes over a hundred iterations with significant *in vivo* DE genes (grey) and the intersection of the top 200 SGS-responsive genes with significant *in vivo* DE genes (orange). (e) Overlap between the top 200 SGS-responsive genes inferred from the WT sample and significant *in vivo* DE genes (*FDR* < 0.05, |Log2(FC)| > 0.25). (f) Overlap between significant GO terms inferred from the top 200 SGS-responsive genes and significant GO terms inferred from the top 200 *in vivo* DE genes. (g) Significantly enriched GO biological processes terms using the top 200 SGS-responsive and their corresponding associated genes.

The chemokine (C–C motif) receptor 2 (*Ccr2*) gene plays a complex and multifaceted role in glioblastoma, with both pro-tumor and anti-tumor effects depending on the cell type and context ^17^. The *Ccr2* receptor is located mainly on the surface of tumor-associated macrophages (TAM) ^18,19^. Since monocytes are precursors to macrophages and dendritic cells in the myeloid cell lineage ^20^, we were interested in studying the functional importance of *Ccr2* in monocytes. We isolated monocytes from the scRNA-seq data to study *Ccr2* function (**Supplementary Fig. 1a**). A total of 3048 WT monocytes were used as input for scSGS analysis (**Fig. 2c**). Then, using the Spline-HVG algorithm, we tested for high variability in *Ccr2* expression. *Ccr2* was identified as highly variable in the dataset (2269 *Ccr2+* and 779 *Ccr2-* cells), making it a good candidate for scSGS analysis (**Supplementary Table 13**). We then performed the scSGS analysis and identified 491 significant SGS-responsive genes (*FDR* < 0.01). The SGS-responsive genes were then ranked based on increasing P-value. We identified 460 DE genes by comparing the *in vivo* KO and WT monocytes. From the top 200 SGS-responsive genes, 72 genes were also significant DE genes (**Fig. 2e**). *Ccr2* was ranked the top SGS-responsive gene, followed by Glycoprotein Nmb (*Gpnmb*), a gene mainly expressed in microglia and macrophages, that plays an important role in the regulation of immune/inflammatory responses ^21^. The gene ranked third was Cathepsin B (*Ctsb*), a gene linked to immune cell infiltration and immunosuppression in gliomas ^22^.

The SGS-responsive genes were significantly biologically connected, as shown by the STRING interaction network (**Fig. 2b** and **Supplementary Table 12**, *P* < 1.0e-16). The links in STRING interaction networks represent functional associations between genes from text mining and experimental reports. These associations include both direct and indirect interactions between genes or their products. Thus, our results suggest abundant functional connectivity between *Ccr2* and the SGS-responsive genes. To validate that SGS-responsive genes are biologically significant, we iteratively randomly selected 200 genes from the dataset and checked their overlap with *in vivo* DE genes (Mean overlap = 6.82). The overlap between the top 200 SGS-responsive genes and the *in vivo* significant DE genes was higher (72 genes) (**Fig. 2d**).

To further analyze the effect of the cell-cycle stage on the scSGS analysis, we split the monocyte population into the G1 phase and S phase monocytes using the ‘*CellCycleScoring’* function from Seurat v5.0.1. We identified 126 and 133 SGS-responsive genes (*FDR* < 0.01) from the G1 and S phase subsets, respectively. Among the common SGS-responsive genes from the whole WT monocyte sample, G1 phase and S phase cells; 59 genes were SGS-responsive in the G1 phase only, and 66 genes were SGS-responsive in the S phase only. Among all three groups, 67 genes were commonly SGS-responsive (**Supplementary Fig. 2a**). This suggests that scSGS analysis is sensitive to cell states (cell cycle in this case) and identifies state-specific responsive genes.

Next, to infer the function of *Ccr2* we conducted gene set enrichment analysis with the GO biological processes ontology database (**Fig. 2g and Supplementary Table 3)**. We chose the top 200 SGS-responsive genes for enrichment analysis (**Supplementary Table 2**). The enrichment analysis identified 293 significantly enriched pathways (*FDR* < 0.05), out of which 109 pathways were also found to be enriched using *in vivo* DE genes (**Fig. 2f**). The enrichment analysis revealed 5 genes associated with the *Regulation of Macrophage Activation*. The study by Pombo Antunes *et al*. and other past studies have shown the importance of the *Ccr2*/*Ccl2* pathway in tumor-associated macrophage activation and influx into the tumor microenvironment, which corresponds with our finding ^23,24^. Furthermore, the knockout of *Ccr2* resulted in a steep decline in monocytes and monocyte-derived tumor-associated macrophage (TAM 1) in the tumor microenvironment (**Supplementary Fig. 2b)**. The *in vivo* KO study also showed an overall increase of T cells and B cells in the tumor microenvironment, which corresponds with our finding of *Negative Regulation of Leukocyte Activation* as an enriched ontology. Thus, the functional predictions made by scSGS were validated by the *in vivo Ccr2* KO study by Pombo Antunes *et al*. The enrichment analysis also identified three genes associated with the *Regulation of Dendritic Cell Differentiation* which corresponds with a study by Chiu *et al*. ^25^ showing impaired lung dendritic cell activation specifically in *Ccr2* knockout mice. Since *Ccr2* is associated with monocyte differentiation and chemotaxis, *Positive Regulation of Mononuclear Cell Migration* was also captured by scSGS analysis ^26^.

Therefore, scSGS successfully identified *Ccr2*-related functions from the significant SGS-responsive genes. We further demonstrated that the inferred genes were functionally connected and, more importantly, the predicted functions were consistent with those reported in *Ccr2* studies.

### Real data scSGS analysis unveils *Kdm6b* function in embryonic mice neurons

Lysine Demethylase 6B (*Kdm6b*, aka *Jmjd3*), is a key agent responsible for removing the histone mark histone H3-lysine27 tri-methylation, inherently playing a crucial role in transcriptional activation ^27^. *Kdm6b* is also actively involved in cell differentiation and organ development, neural stem cell generation, and neurogenesis ^28,29^. A study by Wang *et al*. ^30^, comprehensively studied the role of *Kdm6b* in motor neuron development in the mouse spine on a single-cell level. They reported that during spinal cord development, *Kdm6b* is crucial for the diversification of motor neurons to distinct subtypes corresponding to different muscle targets. We collected our second scRNA-seq dataset from this study of motor neurons from WT and *Kdm6b*^-*/*-^ conditional KO mice (**Fig. 3a**).

**Fig. 3.**
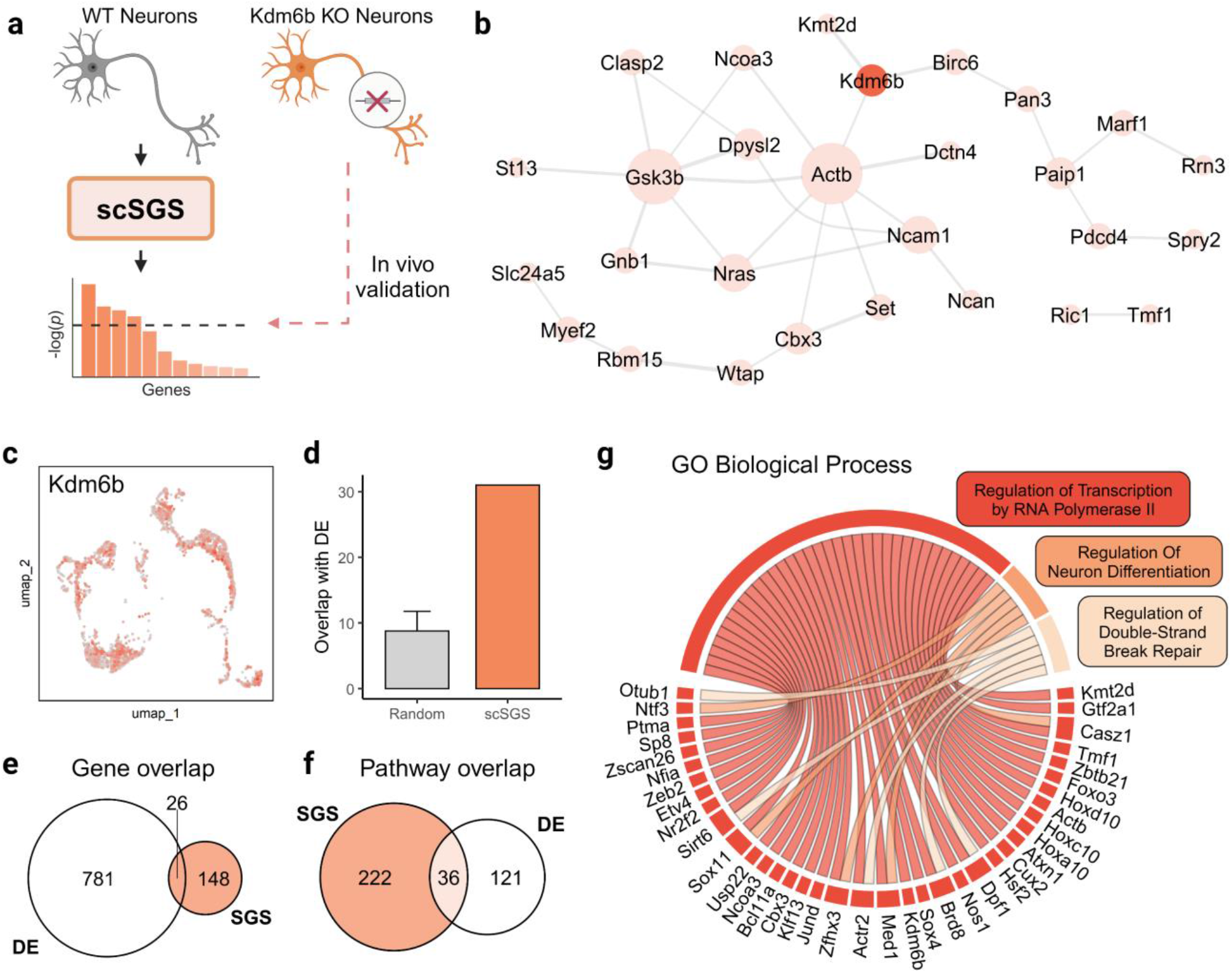
SGS-responsive genes inferred from *Kdm6b* perturbation. (a) Experimental design of scSGS analysis on WT and *Kdm6b*-KO motor neurons from mouse embryonic spine. (b) STRING interaction network of top 50 SGS-responsive genes. Edge thickness indicates the strength of data support and node size indicates node degree. (c) *Kdm6b* expression in WT MN lineage neurons only. (d) The intersection of 200 randomly selected genes over a hundred iterations with significant *in vivo* DE genes (grey) and the intersection of the top 200 SGS-responsive genes with significant *in vivo* DE genes (orange). (e) Overlap between the 174 SGS-responsive genes inferred from the WT sample and significant DE genes (*FDR* < 0.05, |Log2(FC)| > 0.25) from *in vivo* experiment. (f) Overlap between significant GO terms inferred from the SGS-responsive genes and significant GO terms inferred from the top 200 *in vivo* DE genes. (g) Significantly enriched GO biological processes terms using the SGS-responsive genes and their corresponding associated genes.

For this dataset, the motor neuron (MN) lineage (*Slc18a3+* neurons) cells were selected to mimic the original study (**Supplementary Fig. 1b**). After preprocessing and filtering the scRNA-seq data, we extracted the expression profiles of only the 3,156 WT cells for scSGS analysis (**Fig. 3c**). The *Kdm6b* gene expression was identified as highly variable in the WT MN lineage neurons (1650 *Kdm6b+* and 1506 *Kdm6b-* cells) (**Supplementary Table 13**). The analysis identified 174 SGS-responsive genes (*FDR* < 0.01) (**Supplementary Table 4**). The SGS-responsive genes were then ranked based on increasing P-value. From the 174 SGS-responsive genes, 26 genes were also significant DE genes when comparing the *in vivo* KO and WT samples (**Fig. 3e**). The target gene, *Kdm6b*, topped the list. G Protein Subunit Beta 1 (*Gnb1*) was ranked second. Past studies have shown that mutations in the *Gnb1* gene are associated with neuronal developmental delay and epilepsy ^31^. Dihydropyrimidinase-like 2 (*Dpysl2*), a known regulator of neural stem cell differentiation in rats ^32^, ranked fourth. *Dpysl2* is also associated with neurodevelopmental disorders such as autism spectrum disorders and intellectual disability in mice ^33^. Chromobox homolog 3 (*Cbx3*), which ranked fifth, plays a key role in maintaining lineage specificity in the neural differentiation process ^34^. The general trend captured by scSGS analysis suggests an association of *Kdmb6* with neural differentiation and development, which is also the conclusion of the *in vivo* KO study by Wang *et al*.

The top 50 SGS-responsive genes were significantly biologically connected, as shown by the STRING interaction network (**Fig. 3b** and **Supplementary Table 12**, *P* < 0.0038). *Birc6, Actb*, and *Kmt2d* had direct links with the *Kdm6b* gene. Additionally, to validate that scSGS analysis detects biologically significant genes, we iteratively randomly selected 200 genes from the dataset and checked their overlap with *in vivo* DE genes (Mean overlap = 10.74). The SGS-responsive genes had a higher overlap with the *in vivo* DE results (31 genes) (**Fig. 3d**).

Next, to infer the biological processes affected by the SGS pattern of *Kdm6b* we conducted gene set enrichment analysis using all 174 SGS-responsive genes (**Supplementary Table 4)** with the GO biological processes ontology database (**Fig. 3g** and **Supplementary Table 5**). scSGS identified 258 significantly enriched pathways (*FDR* < 0.05), out of which 36 pathways were also enriched by the top 200 *in vivo* DE genes (**Fig. 3f**). *Positive Regulation of Transcription by RNA Polymerase II* was the most significant ontology term which is consistent with a study by Estarás *et al*. ^35^, which shows that *Kdm6b* drives RNA polymerase II progression through H3K27me3-enriched gene bodies. A study by Williams *et al*. ^36^ showed increased JMJD3, the protein coded by *Kdm6b*, levels in response to ionizing radiation-induced DNA double-stranded breaks, which is consistent with our finding of *Regulation of Double-Strand Break Repair* as an enriched ontology. Finally, *Regulation of Neuron Differentiation* was also an enriched term, which is consistent with the *in vivo* KO study by Wang *et al* ^30^.

Therefore, with regard to the significant SGS-responsive gene, our results successfully captured *Kdm6b*-related functions reported in the original study from the significant SGS-responsive genes. We also could predict unreported *Kdm6b-*related functions using just the WT cells, and more importantly, the predicted functions were consistent with those found in other *Kdm6b* KO studies.

### The scSGS tool is robust and scalable

Because of the large heterogeneity in biological samples and a plethora of experimental biases, the reproducibility and robustness of algorithms for similar experimental conditions are important. To test the scSGS analysis framework, we selected three human PBMC datasets of varied sizes from the 10x Genomics datasets repository (**Fig. 4a**). All three datasets were pre-processed, filtered, and annotated using the same parameters and metrics. We selected CD4+ T cells from each dataset (Number of CD4+ T cells: PBMC 5K – 2480 cells, PBMC 10K – 4980 cells, and PBMC 20K – 10,170 cells) for scSGS analysis (**Supplementary Fig. 1 c, d and e**). Then, using the Spline-HVG method, we identified *STAT1* as a common highly variable gene in the CD4+ T cells samples (**Supplementary Table 13**). Because of the extensive literature available on the *STAT1* gene function in CD4+ T cells, we chose the *STAT1* gene for testing ^37-39^. The scSGS analysis with *STAT1* as the target gene revealed 410, 470, and 902 significant SGS-responsive genes (*FDR* < 0.01) in the PBMC 5K, PBMC 10K, and PBMC20K datasets respectively (**Supplementary Tables 8, 9, and 10**). A total of 49 genes were commonly identified as SGS-responsive across all three datasets (**Fig. 4d**). The PBMC 10K and PBMC 20K datasets overlapped considerably with 219 common SGS-responsive genes. scSGS identified 5, 88, and 70 significantly enriched pathways (*FDR* < 0.05) in the PBMC 5K, PBMC 10K, and PBMC 20K datasets, respectively. Five pathways were common across all datasets (**Fig. 4e**). All the pathways enriched for the PBMC 5K dataset were also enriched in the PBMC 10K and PBMC 20K datasets. The PBMC 10K and the PBMC 20K datasets had 53 common pathways.

**Fig. 4.**
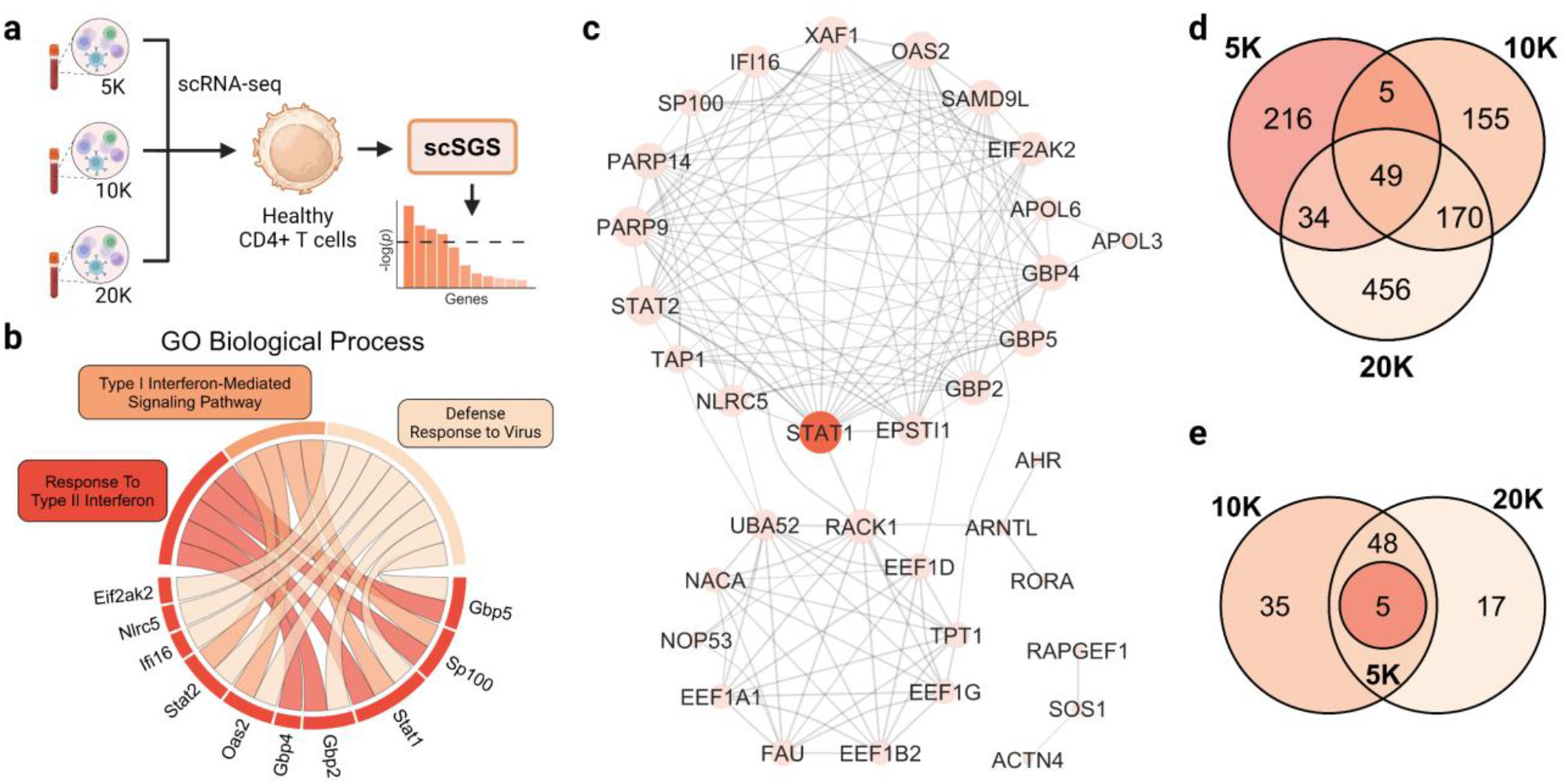
scSGS reproduces similar *STAT1* functional inferences from three independent PBMC datasets. (a) Extractions of CD4+ T cells from three independent Human PBMC datasets (PBMC 5K, PBMC 10K, PBMC 20K) for scSGS analysis. (b) Significantly enriched GO biological processes terms using the common SGS-responsive genes from the three PBMC datasets. (c) STRING interaction network of common *STAT1*-specific SGS-responsive genes from the 3 datasets. Edge thickness indicates the strength of data support and node size indicates node degree. (d) Overlap of significant SGS-responsive genes from the three PBMC datasets. (e) Overlap of significant GO terms inferred from the top 200 SGS-responsive genes from the three PBMC datasets, respectively.

The high overlap among the enriched pathways implies similar functional inferences across the datasets, proving that scSGS is a robust framework. Furthermore, similar functional inferences from the three datasets with varied sample sizes prove that the scSGS tool is scalable.

To assess the replicability of results within the same dataset, we iteratively randomly split the CD4+ T cell population from the PBMC20K dataset into two equal cell subsets and performed scSGS analysis using STAT1 as the target gene (**Supplementary Fig. 5**). Minor shifts were observed in the FDR values of common SGS-responsive genes between splits; however, the rank order of gene outputs remained significantly correlated (P-value < 0.05). The average log2(FC) values across the two splits were also perfectly correlated in each iteration. These findings demonstrate that gene silencing patterns for the target gene are stable within the dataset for a particular cell type, supporting the robustness of the scSGS method.

**Fig. 5.**
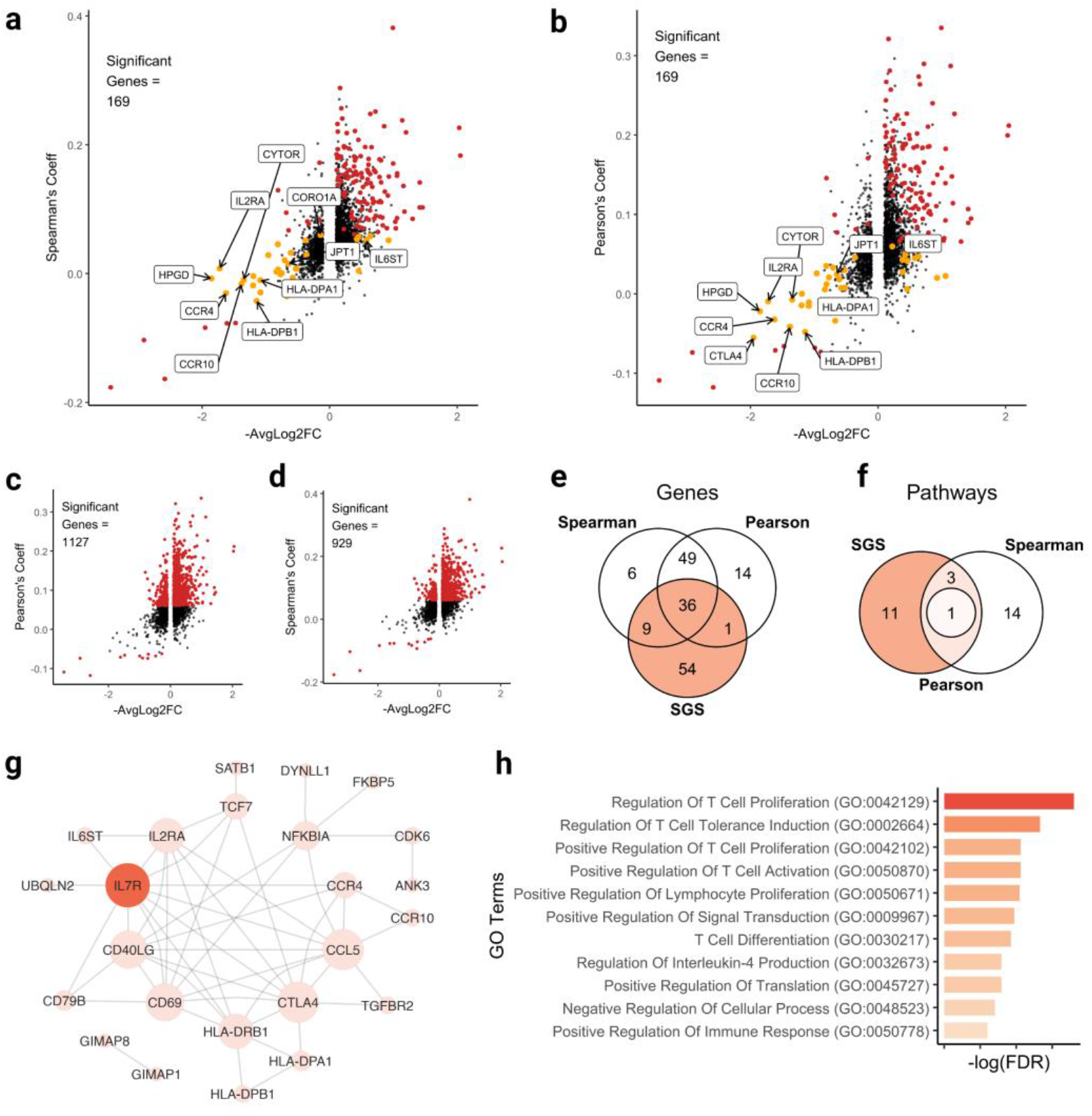
scSGS identified genes associated with *IL7R* overlooked by correlation analysis. (a) Relationship between Spearman’s rank correlation coefficient and scSGS average Log2(FC) (orange - exclusive scSGS identified, red - SGS-responsive genes that are also significantly correlated by Spearman’s correlation test). (b) Relationship between Pearson’s correlation coefficient and scSGS average Log2(FC) (orange - exclusive scSGS identified, red - SGS-responsive gene that is also significantly correlated by Pearson’s correlation test). (c) Significantly correlated genes identified by Spearman’s correlation test are highlighted in red. (d) Significantly correlated genes identified by Pearson’s correlation test are highlighted in red. (e) Overlap between top 100 SGS-responsive, top 100 correlated genes by Spearman’s correlation test and top 100 correlated genes by Pearson’s correlation test. (f) Overlap between significant GO terms inferred from top 100 SGS-responsive, top 100 correlated genes by Spearman’s correlation test and top 100 correlated genes by Pearson’s correlation test. (g) STRING interaction network of the 54 unique SGS-responsive genes with *IL7R*. Node size indicates node degree. (f) Significant GO Terms uniquely identified by SGS-responsive genes.

Gene set enrichment analysis was performed on the 49 common SGS-responsive genes with the GO Biological Process database. The SGS-responsive genes were enriched with genes associated with the *Defense Response to Virus, Response to Type II Interferon*, and *Type I Interferon-Mediated Signaling Pathway* which are all known *STAT1*-associated ontologies ^39-41^ (**Fig. 4b** and **Supplementary Table 11**).

In addition, the SGS-responsive genes were found to be biologically connected, with most of them having direct interaction links with the *STAT1* gene, as shown by the STRING interaction network (**Fig. 4c** and **Supplementary Table 12**, *P* < 1.0e-16). The scaled expressions of 18 genes directly linked to *STAT1* (*TAP1, XAF1, SAMD9L, APOL6, RACK1, EIF2AK2, EPSTI1, IFI16, OAS2, NLRC5, GBP4, GBP2, GBP5, PARP14, PARP9, SP100* and *STAT2*) were plotted as dot plots (**Supplementary Fig. 3**). Past studies have shown that the *SP100* gene is associated with the potential to restrict the initial stages of viral infection ^42^ and plays a role in the type II interferon (IFN-γ) signaling pathway ^43^. Both of these pathways are also associated with the *STAT1* gene ^44^. The *PARP14* and *PARP9* are known cross-regulators of the *STAT1* gene and control macrophage activation through IFN-γ signaling ^45^. Several past studies have shown a link in *GBP5* expression with *STAT1* expression in viral infections through the JAK-STAT pathway ^46,47^. We also observed similar expression patterns for these genes across the three datasets, suggesting that scSGS analysis results can be reproduced for similar cell types from the same tissue across independent datasets. To check for experimental or biological biases, we further randomly selected 18 SGS-non-responsive genes and observed their gene expression patterns (**Supplementary Fig. 4**). No apparent trend or difference in expression pattern was consistently observed across the three datasets.

Thus, we have systematically demonstrated that scSGS analysis is robust and can reproduce similar results, i.e., infer similar SGS-responsive genes and enriched gene functions for similar experimental conditions from independent datasets. Our results also could accurately capture *STAT1*-related functions using the SGS-responsive genes.

### scSGS and correlation are not correlated

Next, we set out to determine if SGS-responsive genes are similar to genes correlated with the target gene using Spearman’s and Pearson’s correlation. We used the PBMC20K dataset and extracted only the CD4+ T cells (10,170 cells) (**Supplementary Fig. 1e**). We filtered out genes with less variable signals, preserving only the top 5000 HVGS for analysis. This allowed for a more equitable comparison of results against correlation metrics that are sensitive to data sparsity. We selected *IL7R* as the target gene, as it was an HVG identified by Spline-HVG (**Supplementary Table 13**) and due to extensive literature available on *IL7R* function in CD4+ T cells ^48,49^. The scSGS analysis with *IL7R* as the target gene revealed 169 significant SGS-responsive genes (*FDR* < 0.01) (**Supplementary Table 6**). We also performed Spearman’s asymptotic t-test and Pearson’s product-moment test to identify genes significantly correlated with *IL7R*. By using an FDR cutoff of < 0.01, we identified 1127 significant genes from Pearson’s correlation test (**Fig. 5c**) and 929 significant genes from Spearman’s correlation test (**Fig. 5d**).

To investigate the biological significance of SGS-responsive genes, we focused on the top 10 genes significantly identified by SGS but not by Spearman’s and Pearson’s correlation metrics (**Fig. 5a, b**). Among these, cytoskeletal regulatory long non-coding RNA, *CYTOR*, exhibits increased expression levels following T cell activation, a process strongly influenced by IL7R signaling ^50^. Additionally, *IL2RA* (Interleukin-2 Receptor Subunit Alpha), a key regulator of immunity, is expressed on regulatory T cells (Tregs), whose development and maintenance are dependent on IL7R signaling ^51^. The *IL6ST* gene, encoding a signal transducer for the interleukin-6 (IL-6) cytokine, has also been linked to *IL7R* gene polymorphisms, which are associated with increased susceptibility to smoking-related asthma ^52^. Moreover, a study reported an increase in HLA-DP+ cells within the CD127+ (*IL7R*) T cell population ^53^. Furthermore, *CCR4* expression has been found to be elevated in CD127-CD4+ T cells ^54^, highlighting the differential immune responses mediated by IL7R signaling. These findings highlight the ability of scSGS to identify key *IL7R-related* genes that were not detected by traditional correlation metrics, demonstrating its sensitivity in uncovering biologically significant genes.

Both correlation metrics identified a substantially higher number of significant peaks, suggesting an inflated rate of false positives. In contrast, scSGS detected a smaller set of significant genes, but these were more likely to be biologically relevant, indicating improved statistical power.

In addition, the SGS-responsive genes were found to be biologically connected, with most of them having direct interaction links with the *IL7R* gene, as shown by the STRING interaction network (**Fig. 5g** and **Supplementary Table 12**, *P* < 1.17e-12). We conducted gene set enrichment analysis on the top 100 SGS-responsive genes and compared these results to those obtained from the top 100 genes identified using Pearson’s and Spearman’s correlation tests. Notably, 54 of the SGS-responsive genes were not among the top 100 genes ranked by either correlation metric (**Fig. 5e**). We also found 11 significantly enriched (FDR < 0.05) GO Biological Process terms that were exclusively identified by the top 100 SGS-responsive genes (**Fig. 5f**). The SGS-responsive genes were enriched with known CD4+ T cell-specific pathways affected by *IL7R*: *Regulation of T Cell Proliferation* ^55^, *Positive regulation of T Cell Activation* ^48^, *T Cell Differentiation*^*56*^, *Regulation of Interleukin-4 Production* ^57^(**Fig. 5h** and **Supplementary Table 11**).

Therefore, our results demonstrate that scSGS fundamentally differs from Spearman’s rank correlation and Pearson’s product-moment correlation. We could also predict known, *IL7R*-specific, T cell activation and differentiation functions using just the healthy PBMC dataset. These results highlight the ability of scSGS to uncover gene relationships, often overlooked by correlation analysis, within single-cell data and its potential to advance our understanding of cellular processes.

## Discussion

In this study, we showcased the purpose and functionality of studying stochastic gene expression using the scSGS framework. We evaluated the inference power of the framework using two *in vivo* gene perturbation studies and 3 healthy scRNA-seq datasets. The SGS-responsive genes showed high overlap with differentially expressed genes from *in vivo* studies. Moreover, scSGS identified genes that had similar stochastic silencing patterns with the gene of interest. Then, we evaluated the robustness and scalability of the framework using three independent peripheral blood mononuclear cell (PBMC) datasets with varied cell numbers. The functional inferences across all three datasets were consistent, and a higher number of SGS-responsive genes and pathways were identified as the number of cells used for analysis increased. Using the PBMC dataset with the largest number of cells, we compared scSGS against two correlation metrics and were able to uncover functionally relevant genes that were not detected by correlation analysis.

Our main goal in this work was to introduce a way to study gene function using purely observational statistics. Past studies have associated dropouts in scRNA-seq gene expression with biological function and phenotype. These studies showcase the potential of using scRNA-seq dropouts for various data analysis procedures, including differential expression ^58,59^, feature selection ^12^, and clustering of cell types by dropout patterns ^11^. To the best of our knowledge, the scSGS framework is the first cell type-specific virtual gene function inference tool based on the silencing pattern of scRNA-seq data. Several gene regulatory network-based virtual knockout tools have been developed for studying perturbation effects using WT scRNA-seq data. They use gene-gene co-expression graphs from WT scRNA-seq data as a model and study the effects of gene perturbations using machine-learning techniques such as manifold alignment ^60^, signal propagation ^61^, and variational graph autoencoders ^62^. These methods, however, require accurate computation of gene regulatory networks, which is still a difficult computational problem in single-cell genomics. Gene regulatory network computation is also computationally intensive resulting in high runtimes (**Supplementary Table 14**). We were able to get similar power as the graph manifold alignment-based scTenifoldKnk at significantly lower runtimes (**Supplementary Fig. 6**). Moreover, the undirected graph-based inputs of these computational techniques cannot capture the directional change (up-regulation or down-regulation) in gene expression caused by perturbations. The scSGS framework, relying on the stochastic silencing patterns of genes, generates a fold change statistic which signifies the directional change of the SGS-responsive gene expressions.

Pearson’s correlation is a widely used correlation metric for studying gene co-expression patterns. However, it assumes that the data follows a linear relationship and that both variables are normally distributed, making it unsuitable for analyzing gene expression data where such assumptions are often not met. In contrast, Spearman’s correlation, which ranks data rather than assuming a specific distribution, is better suited. Nonetheless, Spearman’s correlation test with asymptotic t approximation has several limitations in biological studies. It is less sensitive to non-monotonic or complex nonlinear relationships, common in gene interactions. Additionally, the approximation struggles with data that contain ties, i.e., when multiple cells have identical gene expression levels or ranks. The presence of ties can distort the rank-based Spearman correlation and reduce the accuracy of the t approximation. This makes the test less robust in the face of non-ideal data distributions which are typical in biological datasets. These limitations highlight the need for additional methods, like scSGS, to capture biologically relevant associations more effectively.

The stochastic silencing of genes in some cells can be imagined as miniature naturally occurring perturbation studies. Our analysis framework aimed to utilize this naturally occurring silencing in transcription patterns to infer gene function. Even though we have used DE results from *in vivo* gene perturbations to validate the results from scSGS, SGS-responsive genes are not completely comparable to DE genes from knockout experiments, because of survivorship bias. Hence, we validated most of our findings with prior knowledge and a literature search. However, like any other computational tool, the scSGS tool has its shortcoming. The drawbacks of the scSGS framework arise from the fact that it is based solely on observational statistics. The Wilcoxon rank sum test can have reduced statistical power when sample sizes are small. This can make it challenging to detect true differences between groups. Therefore, the inference power and accuracy of the scSGS framework are directly proportional to the sample size, i.e., the number of single cells in the study. Similarly, even in large groups of cells if the target gene has a significantly high or low number of cells expressing it, splitting the cell sample based on the binarized expression of the genes is disproportional, leaving a small number of cells on either the active or silenced sample. In the PBMC datasets with different cell numbers, we observed more genes and pathways associated with *STAT1* with the increase in the number of cells present in the dataset. Additionally, the scSGS analysis is prone to selection bias, i.e., using a sample of cells for analysis rather than the whole population will lead to slightly different inferences. To control this bias, the analysis was conducted in a cell state-specific manner using only a single, biologically validated, cell type or cell-cycle phase.

Overall, the scSGS tool effectively utilizes the individual gene expression patterns to infer gene function from large multidimensional scRNA-seq data. The silencing patterns created by transcriptional bursting and individual gene expression heterogeneity captured by scRNA-seq are key to more comprehensive gene function studies. scSGS provides a computationally non-intensive yet robust method for analyzing these patterns. Moreover, the bias-free prediction power of scSGS enables the prediction of gene knockouts, so that the KO gene’s functions can be revealed in a cell-type-specific manner, reducing the need for exploratory animal trials. We anticipate that scSGS will be widely applied in predicting gene function in single-cell biomedical research.

## Materials and Methods

### Preprocessing of publicly available data sets

We used publicly available scRNA-seq data sets. The specifics and source of the real scRNA-seq datasets used are listed in **Supplementary Table 1** ^16,30,63-65^. We filtered out genes expressed in ≤ 15 cells. We further filtered out cells with >5% mitochondrial read counts and ≤ 500 expressed genes. The top 1% of cells with a large number of reads were excluded to reduce outliers and control the library size. We followed a general pipeline from the Seurat (v5.0.1) package ^66^ to process the data, starting with log normalization of scRNA-seq samples using the ‘*NormalizeData’* function. Mitochondrial and ribosomal genes were excluded from the datasets. We then scaled the data and performed principal component analysis (PCA) using the top variable genes computed by the ‘*FindVariableFeatures’* function. We used the first 40 principal components for dimensionality reduction via uniform manifold approximation and projection (UMAP). Then, we identified cell clusters using the Louvain algorithm via the ‘*FindClusters’* function with a resolution parameter of 0.8. Unless metadata was provided with the dataset, we used the ScType method ^15^ using the cell type markers from the ScType database to automatically annotate the identified cell clusters. Finally, to approximate the cell-cycle stages of cells, we used the ‘*CellCycleScoring’* function using Seurat’s in-built S phase and G2/M phase gene lists.

## Finding Highly Variable Genes using 3D spline interpolation – Spline-HVG

We filtered out genes with zero mean expression and identified highly variable genes (HVGs) using a 3D spline curve-based method developed by Cai *et al*. ^67^. For the scRNA-seq expression over the counts *K*_*ij*_, across all genes *i* = 1, …, *m* and cells *j* = 1, …, *n*, we computed three gene summary statistics: log of mean expression (*logµ*), log of coefficient of variance (*logCV*), and dropout rate (*DR*). The mean and CV for each gene were computed across all cells and then *log*(*x* + 1) transformed.

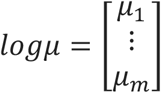

Where for *i* ∈{1,…,*m*},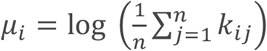

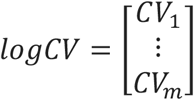

Where for *i* ∈{1,…,*m*},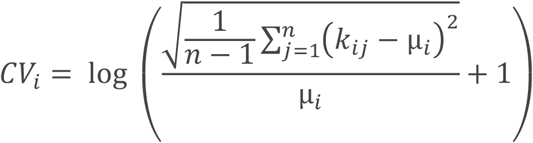

We then computed the dropout rate as the fraction of cells with zero expression for the given gene.

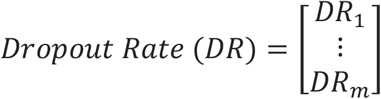

Where for *i* ∈{1,…,*m*},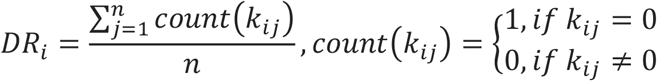

We characterized every gene by these three variables, which in turn defined its unique position in the 3D space. We sorted the genes in ascending order of dropout rate. To capture the accumulated variability, we calculated the cumulative sum of the square root of squared differences.

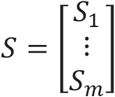

Where for *i* ∈{1,…,*m*},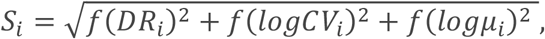

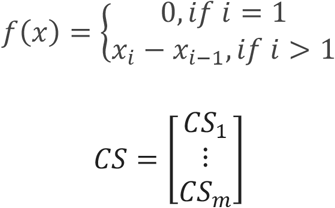

Where for *i* ∈{1,…,*m*},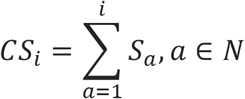

We performed cubic spline fitting for each variable (*logµ, logCV, DR*) with *CS* using the *‘smooth*.*spline’* function from the R stats package. We used 15 degrees of freedom and set the smoothing parameter to 0.75 to control and balance the smoothness, mimicking the MATLAB-based *‘SPLINEFIT’* function ^68^. This function handles noisy data and removes unwanted oscillations in the spline curve. Values were predicted for each gene at points along the cumulative sum (*CS*) axis using the fitted splines. The shortest distance, d, from each data point to the spline curve was computed using the *‘matchpt’* function from the Biobase R package ^69^. This distance was used as the measure of the corresponding gene’s variability. Genes with large d were called highly variable.

### Identification of target scSGS genes

Using the top 500 HVGs from the Spline-HVG results, we then selected genes with dropout rates > 0.25 and < 0.75. We used this dropout rate so that, post-classification, each subset had enough cells (at least 25% percent of the total cell population) for statistical testing. We further checked whether the genes identified were transcription factors by searching in a comprehensive list curated by Ng *et al*. ^70^.

### Classification of WT scRNA-seq sample into Active and Silenced cells

To use the count expression as a classifier, we binarized the expression pattern of the target gene (*G*_*j*_) across all cells *j =* 1, …, *n*.

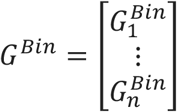

Where for *j* ∈{1,…,*n*},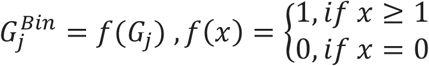

Using the binarized expression 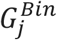, we split the cell population into two subsets: Silenced

(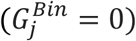) and Active (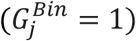).

### Testing for significance of SGS-responsive genes

We compared the silenced sample with the active sample using a fast implementation of the non-parametric Wilcoxon rank-sum test via the Presto package ^71^. We also excluded genes coding for ribosomal protein and mitochondrial genes. We also excluded genes expressed in < 10% of the cells of either group or having an absolute Log2(fold-change) of < 0.1 from testing. Finally, we corrected the P-values using the Benjamini-Hochberg FDR adjustment to obtain adjusted P-values. Genes with an adjusted P-value (*FDR*) of < 0.01 were considered SGS-responsive. To examine the inference power of the scSGS analysis, we computed the intersection of the top 200 scSGS-responsive genes with DE genes from the *in vivo* KO study. We also computed the intersection of 200 randomly chosen genes with DE genes from the *in vivo* KO study. These two intersections were then compared. The randomly chosen non-responsive genes were computed over 100 iterations with a unique random seed on each iteration.

### Gene functional enrichment and network analysis

To identify biological processes and pathways associated with the SGS-responsive or *in vivo* differently expressed genes, we performed gene set enrichment analysis (GSEA) with the KEGG and GO Biological Processes library using Enrichr ^72,73^. We used the top n significant genes (n ≤ 200) with the default settings to run the analysis. We focused on gene sets with significant enrichment (*FDR* < 0.05) and biological relevance to the respective study. To further explore the known gene-gene relationships, we used the top 50 scSGS-responsive genes for protein-protein interaction network analysis using the STRING Database v12 ^74^. We chose a medium interaction confidence score threshold to filter the network and removed isolated nodes, aiming for a balance between inclusivity and reliability. We then performed enrichment analysis for pathway terms to identify significantly overrepresented functional categories within the network.

### Benchmarking

We selected another unsupervised R-based single-cell gene function inference tool, scTenifoldKnk ^60^. scTenifoldKnk like scSGS required only WT gene expression data for predicting virtual gene knockout-affected genes. The benchmarking was performed on the WT monocytes in the *Ccr2* KO dataset from mice glioblastoma. We selected genes using the Spline-HVG algorithm and the cells using random sampling without replacement to create four test datasets. The 5 test datasets were of the following dimensions: test set 1 (1000 genes, 500 cells), test set 2 (1000 genes, 1000 cells), test set 3 (3000 genes, 1000 cells), test set 4 (5000 genes, 1000 cells) and test set 5 (5000 genes, 3000 cells). We ran both scTenifoldKnk and scSGS to evaluate the runtimes and overlaps within their results. The evaluations were implemented on equivalent hardware, composed of a 14-core Intel Core i5-13500 CPU processor at 2.50 GHz with 31.7 GB available random-access memory (RAM). We also compared the analysis results with the real DE analysis using the same 5000 HVGs comparing the experimental WT and KO samples.

### Correlation testing

We conducted correlation analyses to evaluate the relationships between gene expression levels and our target gene. Pearson’s product-moment and Spearman’s rank correlation coefficients were calculated using the ‘cor.test’ function in R (v4.4.1). We used Pearson’s correlation to assess linear relationships, and Spearman’s correlation to capture monotonic, rank-based relationships. For Pearson’s correlation, we computed test statistics from an asymptotic confidence interval based on Fisher’s Z transform. For Spearman’s correlation, we computed P-values via the asymptotic t approximation. To account for multiple comparisons, we adjusted P-values for both Pearson’s and Spearman’s correlations using the Benjamini-Hochberg method to control the false discovery rate (FDR). The adjusted P-values below 0.01 were considered statistically significant.

### Differential gene expression analysis

We employed the ‘*FindMarkers’* function from Seurat (v5.0.1) to conduct differential expression analysis. We used the Wilcoxon rank-sum test with P-value correction for multiple tests using the Benjamini–Hochberg procedure. To identify significant DE genes, we selected genes with *FDR* < 0.05 and |Log2(FC)| > 0.25.

## Data Availability

The sources of data sets underlying this article can be found in **Supplementary Table 1**. No new data were generated in support of this research. An R implementation of the scSGS framework is available at https://github.com/Xenon8778/scSGS.

## Author Contributions

**S.G**.: Conceptualization, Methodology, Visualization, Software, Formal Analysis, Writing - Original Draft. **J.J.C**: Conceptualization, Methodology, Supervision, Writing - Review & Editing, Resources.

## Acknowledgements

We would like to express our appreciation to Dr. Barbara Gastel for her valuable literary support and feedback on manuscript writing.

## Supplementary Figure Legends

**Supplementary Fig. 1**. UMAP plots highlighting cell types/states used for scSGS analysis from the respective datasets.

**Supplementary Fig. 2**. a) Overlap of SGS-responsive genes identified from all WT monocytes, G1 phase monocytes, and S phase monocytes. b) Cell percentages across cell types in WT and *Ccr2* KO samples.

**Supplementary Fig. 3**. Expressions of SGS-responsive genes directly linked with *STAT1* according to STRING interaction analysis in the three PBMC datasets in Active and Silenced subsets.

**Supplementary Fig. 4**. Expressions of 18 randomly selected genes from the three PBMC datasets in Active and Silenced subsets.

**Supplementary Fig. 5**. Relationship between FDR and average Log2(FC) values between two randomly split samples from the same PBM20K dataset over 5 iterations.

**Supplementary Fig. 6**. Overlap of top 100 SGS-responsive and top 100 significant virtual KO genes from scTenifoldKnk with top 100 DE genes from *in vivo Ccr2* KO study

## Supplementary Table Legends

**Supplementary Table 1**. Summary of real scRNA-seq datasets used for scSGS analysis.

**Supplementary Table 2**. Top 200 *Ccr2* specific SGS-responsive genes from WT monocytes in the glioblastoma dataset.

**Supplementary Table 3**. Top 20 significant GO Biological Process terms using top 200 *Ccr2* specific SGS-responsive genes from WT monocytes in the glioblastoma dataset.

**Supplementary Table 4**. *Kdm6b* specific SGS-responsive genes from WT motor neurons in the embryonic mouse spine dataset.

**Supplementary Table 5**. Top 20 significant GO Biological Process terms using *Kdm6b* specific SGS-responsive genes from WT motor neurons in the embryonic mouse spine dataset.

**Supplementary Table 6**. *IL7R* specific SGS-responsive genes from CD4+ T cells in the PBMC20K dataset.

**Supplementary Table 7**. Significant GO Biological Process Terms using the top 100 *IL7R* specific SGS-responsive genes from CD4+ T cells in the PBMC20K dataset.

**Supplementary Table 8**. Top 200 *STAT1* specific SGS-responsive genes from CD4+ T cells in the PBMC5K dataset.

**Supplementary Table 9**. Top 200 *STAT1* specific SGS-responsive genes from CD4+ T cells in the PBMC10K dataset.

**Supplementary Table 10**. Top 200 *STAT1* specific SGS-responsive genes from CD4+ T cells in the PBMC20K dataset.

**Supplementary Table 11**. Top 20 significant pathways obtained from Enrichr after gene set enrichment of common *STAT1* specific SGS-responsive genes from the three PBMC datasets.

**Supplementary Table 12**. Permalinks for STRING interaction analysis using top SGS-responsive genes.

**Supplementary Table 13**. Gene statistics of target genes from respective datasets using the Spline-HVG method.

