## Supplementary Information for "Gene Function Revealed at the Moment of Stochastic Gene Silencing"

### Supplementary Tables

**Supplementary Table 1.** Summary of real scRNA-seq datasets used for scSGS analysis.

| **Data set** | **Experiment Type** | **Organism** | **Cell Type of Interest (# of Cells)** | **Target Gene** | **Source** |
|---|---|---|---|---|---|
| Glioblastoma | scRNA-seq | *Mus musculus* | Monocytes (3048) | *Ccr2* | GSE163120 [1] |
| | | | G1 Phase Monocytes (1241) | | |
| | | | S Phase Monocytes (1157) | | |
| Embryonic spine | scRNA-seq | *Mus musculus* | Neurons (8876) | *Kdm6b* | GSE156609 [2] |
| PBMC – 5k | scRNA-seq | *Homo sapiens* | CD4+ T cells (2480) | *STAT1* | 10x Genomics [3] |
| PBMC – 10k | scRNA-seq | *Homo sapiens* | CD4+ T cells (2543) | *STAT1* | 10x Genomics [4] |
| PBMC – 20k | scRNA-seq | *Homo sapiens* | CD4+ T cells (2830) | *STAT1* | 10x Genomics [5] |

**Supplementary Table 2.** Top 200 *Ccr2* specific SGS-responsive genes from WT monocytes in the glioblastoma dataset.

| Rank | Gene | FDR | Rank | Gene | FDR | Rank | Gene | FDR | Rank | Gene | FDR |
|---|---|---|---|---|---|---|---|---|---|---|---|
| 1 | *Ccr2* | 0.00E+00 | 51 | *Gpx1* | 2.79E-09 | 101 | *Cd74* | 2.16E-06 | 151 | *Ndufa6* | 4.48E-05 |
| 2 | *Gpnmb* | 2.03E-47 | 52 | *Vsir* | 3.38E-09 | 102 | *Timp2* | 2.21E-06 | 152 | *Cdkn2d* | 4.78E-05 |
| 3 | *Ctsb* | 8.71E-28 | 53 | *Tmem176b* | 3.58E-09 | 103 | *Pirb* | 2.43E-06 | 153 | *Ypel3* | 4.87E-05 |
| 4 | *Fth1* | 1.38E-25 | 54 | *Susd3* | 6.81E-09 | 104 | *Fxyd5* | 2.46E-06 | 154 | *Mef2c* | 4.87E-05 |
| 5 | *Ftl1* | 5.45E-25 | 55 | *Cyba* | 7.67E-09 | 105 | *Fnip2* | 2.61E-06 | 155 | *Fcgrt* | 5.77E-05 |
| 6 | *Ms4a6b* | 4.22E-24 | 56 | *Ifngr1* | 1.24E-08 | 106 | *Dbi* | 2.71E-06 | 156 | *Sp100* | 5.81E-05 |
| 7 | *Fabp5* | 2.58E-22 | 57 | *Twf2* | 1.51E-08 | 107 | *AW112010* | 2.71E-06 | 157 | *Capg* | 6.05E-05 |
| 8 | *H2-DMa* | 1.97E-21 | 58 | *Rgs1* | 2.27E-08 | 108 | *Emp1* | 2.81E-06 | 158 | *Bhlhe40* | 6.08E-05 |
| 9 | *Cstb* | 4.74E-20 | 59 | *Rbfa* | 2.82E-08 | 109 | *Gpr137b* | 2.83E-06 | 159 | *Psma1* | 6.93E-05 |
| 10 | *Cd63* | 5.75E-20 | 60 | *Ninj1* | 3.23E-08 | 110 | *Tgfbi* | 3.03E-06 | 160 | *Arhgef6* | 7.10E-05 |
| 11 | *Il7r* | 7.00E-19 | 61 | *Atp5c1* | 3.62E-08 | 111 | *Ly86* | 3.39E-06 | 161 | *Nfkbia* | 8.11E-05 |
| 12 | *H2-DMb1* | 7.00E-19 | 62 | *Arhgdib* | 3.96E-08 | 112 | *Arpc4* | 3.63E-06 | 162 | *Clec12a* | 8.36E-05 |
| 13 | *Pld4* | 1.53E-18 | 63 | *Arpc1b* | 4.53E-08 | 113 | *H2-D1* | 4.52E-06 | 163 | *Tppp3* | 8.36E-05 |
| 14 | *Hspa8* | 4.84E-18 | 64 | *Slc25a5* | 6.94E-08 | 114 | *Gm10116* | 4.86E-06 | 164 | *Plin2* | 8.68E-05 |
| 15 | *Cdk2ap2* | 3.10E-17 | 65 | *Ccl6* | 6.94E-08 | 115 | *Arl6ip5* | 5.44E-06 | 165 | *Eif3h* | 8.68E-05 |
| 16 | *Lgals3* | 4.07E-17 | 66 | *Gnai2* | 1.23E-07 | 116 | *Kctd12* | 5.90E-06 | 166 | *Tm6sf1* | 8.77E-05 |
| 17 | *Actr3* | 6.61E-17 | 67 | *Nr1h3* | 1.72E-07 | 117 | *Pold4* | 7.02E-06 | 167 | *Cd72* | 8.87E-05 |
| 18 | *Coro1a* | 1.06E-16 | 68 | *Slc7a11* | 1.72E-07 | 118 | *Naaa* | 8.34E-06 | 168 | *Uqcrfs1* | 9.95E-05 |
| 19 | *Ms4a6c* | 3.11E-16 | 69 | *Bin2* | 1.82E-07 | 119 | *Ly6a* | 8.34E-06 | 169 | *Ckb* | 9.95E-05 |
| 20 | *Clec4a2* | 5.77E-15 | 70 | *Btg2* | 2.71E-07 | 120 | *Clec4a3* | 8.38E-06 | 170 | *Asah1* | 9.98E-05 |
| 21 | *Ctsd* | 6.08E-15 | 71 | *Cfp* | 2.76E-07 | 121 | *Gpr137b-ps* | 1.03E-05 | 171 | *Aif1* | 1.01E-04 |
| 22 | *Syngr1* | 1.61E-14 | 72 | *Rac2* | 2.98E-07 | 122 | *Tmsb4x* | 1.07E-05 | 172 | *Serinc3* | 1.04E-04 |
| 23 | *Plac8* | 2.25E-14 | 73 | *Clic1* | 3.14E-07 | 123 | *Pilrb2* | 1.07E-05 | 173 | *Ifi30* | 1.08E-04 |
| 24 | *Fcgr3* | 3.15E-14 | 74 | *Capza2* | 3.42E-07 | 124 | *Clec4a1* | 1.07E-05 | 174 | *Crtc3* | 1.10E-04 |
| 25 | *Sirpb1c* | 4.78E-14 | 75 | *Spp1* | 4.15E-07 | 125 | *Ms4a8a* | 1.20E-05 | 175 | *H2-Aa* | 1.15E-04 |

| | | | | | | | | | | | |
|---|---|---|---|---|---|---|---|---|---|---|---|
| 26 | *Cd68* | 1.15E-13 | 76 | *Lamtor4* | 4.30E-07 | 126 | *Cox7a2l* | 1.26E-05 | 176 | *Npc1* | 1.20E-04 |
| 27 | *Cst3* | 1.48E-13 | 77 | *Fcgr2b* | 4.37E-07 | 127 | *Batf3* | 1.34E-05 | 177 | *Selplg* | 1.22E-04 |
| 28 | *Alox5ap* | 4.05E-13 | 78 | *Gm2a* | 4.37E-07 | 128 | *Mycbp2* | 1.34E-05 | 178 | *Camk1d* | 1.25E-04 |
| 29 | *Gdi2* | 6.46E-13 | 79 | *Tkt* | 4.43E-07 | 129 | *Spint1* | 1.35E-05 | 179 | *Triap1* | 1.25E-04 |
| 30 | *Serp1* | 3.95E-12 | 80 | *Pla2g7* | 4.52E-07 | 130 | *Eef2* | 1.61E-05 | 180 | *Ccdc109b* | 1.25E-04 |
| 31 | *Sgk1* | 7.94E-12 | 81 | *Fam111a* | 5.76E-07 | 131 | *Tbxas1* | 1.73E-05 | 181 | *1600014C10Rik* | 1.30E-04 |
| 32 | *Ccr1* | 1.24E-11 | 82 | *Plbd1* | 5.79E-07 | 132 | *Rtcb* | 1.73E-05 | 182 | *Dtnbp1* | 1.40E-04 |
| 33 | *Pld3* | 2.05E-11 | 83 | *Sirpb1b* | 5.98E-07 | 133 | *Atp5b* | 1.93E-05 | 183 | *Acot8* | 1.47E-04 |
| 34 | *Ctsc* | 2.05E-11 | 84 | *Irg1* | 6.62E-07 | 134 | *Lpar6* | 2.05E-05 | 184 | *Uqcrh* | 1.47E-04 |
| 35 | *Erp29* | 5.29E-11 | 85 | *Hp* | 6.92E-07 | 135 | *Ctsz* | 2.19E-05 | 185 | *Pstpip1* | 1.47E-04 |
| 36 | *Ccl9* | 5.40E-11 | 86 | *B2m* | 7.79E-07 | 136 | *Gsn* | 2.22E-05 | 186 | *Exosc5* | 1.53E-04 |
| 37 | *Rassf4* | 6.74E-11 | 87 | *Grn* | 9.04E-07 | 137 | *Anxa5* | 2.24E-05 | 187 | *Arpc3* | 1.53E-04 |
| 38 | *Id2* | 1.22E-10 | 88 | *Ccr5* | 9.22E-07 | 138 | *Basp1* | 2.24E-05 | 188 | *Ppm1m* | 1.53E-04 |
| 39 | *Lair1* | 1.48E-10 | 89 | *Atp6v1g1* | 9.80E-07 | 139 | *Svbp* | 2.45E-05 | 189 | *Polr1d* | 1.56E-04 |
| 40 | *Klra2* | 1.82E-10 | 90 | *Btf3* | 1.04E-06 | 140 | *Clec4d* | 2.72E-05 | 190 | *Slamf9* | 1.57E-04 |
| 41 | *Unc93b1* | 1.82E-10 | 91 | *Gla* | 1.11E-06 | 141 | *Ifi27l2a* | 2.72E-05 | 191 | *Maff* | 1.59E-04 |
| 42 | *Ptpn18* | 3.10E-10 | 92 | *Ighm* | 1.11E-06 | 142 | *Napsa* | 2.85E-05 | 192 | *Pitpna* | 1.70E-04 |
| 43 | *Cd9* | 4.23E-10 | 93 | *Fam20c* | 1.21E-06 | 143 | *Gpi1* | 3.12E-05 | 193 | *Hacd4* | 1.75E-04 |
| 44 | *Cmtm7* | 6.49E-10 | 94 | *Psap* | 1.55E-06 | 144 | *Gas7* | 3.23E-05 | 194 | *Chil3* | 1.79E-04 |
| 45 | *Ifitm2* | 8.12E-10 | 95 | *Dusp22* | 1.70E-06 | 145 | *F11r* | 3.46E-05 | 195 | *Sys1* | 1.83E-04 |
| 46 | *H2afz* | 9.03E-10 | 96 | *Dpysl2* | 1.83E-06 | 146 | *Lilr4b* | 3.46E-05 | 196 | *S100a11* | 1.83E-04 |
| 47 | *Cd52* | 1.01E-09 | 97 | *Trappc1* | 1.84E-06 | 147 | *Ptafr* | 3.60E-05 | 197 | *Ms4a6d* | 1.84E-04 |
| 48 | *Lgals3bp* | 1.63E-09 | 98 | *Pkig* | 1.86E-06 | 148 | *Il1b* | 3.84E-05 | 198 | *Cd84* | 1.90E-04 |
| 49 | *Emb* | 2.53E-09 | 99 | *Ppp1ca* | 2.01E-06 | 149 | *Ucp2* | 3.85E-05 | 199 | *Cd302* | 2.05E-04 |
| 50 | *Fam49b* | 2.63E-09 | 100 | *Slamf7* | 2.01E-06 | 150 | *Ethe1* | 3.97E-05 | 200 | *Eif3i* | 2.05E-04 |

**Supplementary Table 3.** Top 20 significant GO Biological Process terms using top 200 *Ccr2* specific SGS-responsive genes from WT monocytes in the glioblastoma dataset.

| **Term** | **Overlap** | **P-value** | **Adj. P-value** | **Genes** |
|---|---|---|---|---|
| *Phagocytosis (GO:0006909)* | 8/69 | 4.31E-07 | 6.76E-04 | *GSN;BIN2;PLD4;NR1H3;CD302;FCGR2B;AIF1;CORO1A* |
| *Regulation Of Macrophage Activation (GO:0043030)* | 5/31 | 1.31E-05 | 5.23E-03 | *CD74;CD84;NR1H3;FCGR2B;GPR137B* |
| *Positive Regulation Of Mononuclear Cell Migration (GO:0071677)* | 5/31 | 1.31E-05 | 5.23E-03 | *CCR1;LGALS3;AIF1;PLA2G7;CCR2* |
| *Actin Polymerization Or Depolymerization (GO:0008154)* | 6/53 | 1.44E-05 | 5.23E-03 | *PSTPIP1;GSN;TWF2;CAPG;AIF1;GAS7* |
| *Negative Regulation Of Leukocyte Activation (GO:0002695)* | 4/17 | 2.09E-05 | 5.23E-03 | *CD84;GRN;NR1H3;FCGR2B* |
| *Actin Filament Capping (GO:0051693)* | 4/18 | 2.66E-05 | 5.23E-03 | *GSN;CAPZA2;TWF2;CAPG* |
| *Barbed-End Actin Filament Capping (GO:0051016)* | 4/18 | 2.66E-05 | 5.23E-03 | *GSN;CAPZA2;TWF2;CAPG* |
| *Positive Regulation Of Response To External Stimulus (GO:0032103)* | 9/155 | 2.67E-05 | 5.23E-03 | *NFKBIA;GRN;IL1B;NINJ1;LY86;CYBA;PLA2G7;AIF1;CCR2* |
| *Arp2/3 Complex-Mediated Actin Nucleation (GO:0034314)* | 4/19 | 3.34E-05 | 5.57E-03 | *ACTR3;ARPC3;ARPC1B;ARPC4* |
| *Positive Regulation Of Defense Response (GO:0031349)* | 8/124 | 3.55E-05 | 5.57E-03 | *NFKBIA;MEF2C;GRN;IL1B;NINJ1;CYBA;PLA2G7;CCR2* |
| *Actin Nucleation (GO:0045010)* | 4/23 | 7.40E-05 | 1.06E-02 | *ACTR3;ARPC3;ARPC1B;ARPC4* |
| *Regulation Of Dendritic Cell Differentiation (GO:2001198)* | 3/10 | 1.12E-04 | 1.47E-02 | *LGALS3;TMEM176B;FCGR2B* |
| *Regulation Of Peptidase Activity (GO:0052547)* | 4/27 | 1.42E-04 | 1.72E-02 | *CST3;CSTB;SVBP;CTSB* |
| *Phagocytosis, Engulfment (GO:0006911)* | 4/30 | 2.17E-04 | 2.43E-02 | *GSN;BIN2;FCGR2B;AIF1* |
| *Regulation Of Leukocyte Activation (GO:0002694)* | 3/14 | 3.31E-04 | 3.26E-02 | *CD74;CD84;GPR137B* |
| *Positive Regulation Of Phagocytosis (GO:0050766)* | 5/60 | 3.32E-04 | 3.26E-02 | *CAMK1D;IL1B;CYBA;CFP;FCGR2B* |
| *Ceramide Catabolic Process (GO:0046514)* | 3/15 | 4.10E-04 | 3.39E-02 | *ASAH1;GM2A;GLA* |
| *Dendritic Cell Chemotaxis (GO:0002407)* | 3/15 | 4.10E-04 | 3.39E-02 | *CCR1;CCR5;CCR2* |
| *Negative Regulation Of Macrophage Activation (GO:0043031)* | 3/15 | 4.10E-04 | 3.39E-02 | *GRN;NR1H3;FCGR2B* |
| *Positive Chemotaxis (GO:0050918)* | 3/17 | 6.04E-04 | 4.44E-02 | *LGALS3;GPNMB;CORO1A* |

**Supplementary Table 4.** *Kdm6b* specific SGS-responsive genes from WT motor neurons in the embryonic mouse spine dataset.

| Rank | Gene | FDR | Rank | Gene | FDR | Rank | Gene | FDR | Rank | Gene | FDR |
|---|---|---|---|---|---|---|---|---|---|---|---|
| 1 | *Kdm6b* | 0.00E+00 | 51 | *Foxo3* | 5.90E-10 | 101 | *Srsf2* | 1.23E-04 | 151 | *Stmn3* | 4.85E-03 |
| 2 | *Gnb1* | 5.16E-85 | 52 | *4933434E20Rik* | 1.23E-09 | 102 | *Cacng8* | 1.27E-04 | 152 | *Bcl11a* | 4.85E-03 |
| 3 | *Actb* | 1.41E-76 | 53 | *Actr2* | 2.70E-09 | 103 | *Gigyf1* | 1.36E-04 | 153 | *Agpat1* | 4.90E-03 |
| 4 | *Dpysl2* | 3.98E-67 | 54 | *Klhl9* | 3.10E-09 | 104 | *Cask* | 1.45E-04 | 154 | *Ccni* | 5.49E-03 |
| 5 | *Cbx3* | 3.46E-66 | 55 | *Sfxn5* | 4.05E-09 | 105 | *Ubald1* | 1.46E-04 | 155 | *Nudt3* | 5.66E-03 |
| 6 | *Slc24a5* | 5.45E-63 | 56 | *Cramp1l* | 7.65E-09 | 106 | *Mcu* | 1.67E-04 | 156 | *Khdrbs1* | 5.86E-03 |
| 7 | *Sox11* | 9.86E-49 | 57 | *Gm15800* | 1.38E-08 | 107 | *Zbtb44* | 1.85E-04 | 157 | *Sp8* | 5.87E-03 |
| 8 | *Lrrc8a* | 2.20E-48 | 58 | *Upf1* | 3.21E-08 | 108 | *Otub1* | 1.92E-04 | 158 | *Ntng1* | 6.40E-03 |
| 9 | *Ncam1* | 1.09E-41 | 59 | *Rdh13* | 3.24E-08 | 109 | *Spock1* | 2.01E-04 | 159 | *Furin* | 6.40E-03 |
| 10 | *Myef2* | 4.19E-39 | 60 | *Zfp780b* | 5.57E-08 | 110 | *Braf* | 2.95E-04 | 160 | *Smpd3* | 6.40E-03 |
| 11 | *Gsk3b* | 2.59E-38 | 61 | *Sept11* | 6.18E-08 | 111 | *Bmp2k* | 3.46E-04 | 161 | *Sox4* | 6.40E-03 |
| 12 | *Hsf2* | 1.31E-33 | 62 | *Arglu1* | 7.33E-08 | 112 | *Gm38393* | 3.65E-04 | 162 | *Rbfox1* | 6.40E-03 |
| 13 | *Paip1* | 4.91E-30 | 63 | *Slc2a12* | 8.84E-08 | 113 | *Pdzd8* | 3.69E-04 | 163 | *Ntf3* | 6.65E-03 |
| 14 | *St13* | 2.32E-27 | 64 | *Golt1b* | 1.57E-07 | 114 | *Ddhd2* | 4.28E-04 | 164 | *Tmem263* | 7.14E-03 |
| 15 | *Kmt2d* | 7.72E-26 | 65 | *Iqsec1* | 3.27E-07 | 115 | *Hoxc10* | 5.07E-04 | 165 | *F2r* | 7.14E-03 |
| 16 | *Inpp4a* | 2.76E-25 | 66 | *Alkbh5* | 3.91E-07 | 116 | *Lrrc58* | 5.24E-04 | 166 | *Nr2f2* | 7.29E-03 |
| 17 | *Nfia* | 2.19E-24 | 67 | *Pfas* | 4.16E-07 | 117 | *Gtf2a1* | 6.00E-04 | 167 | *Etv4* | 7.65E-03 |
| 18 | *Ncoa3* | 1.04E-23 | 68 | *Igsf3* | 4.65E-07 | 118 | *Tubb5* | 6.39E-04 | 168 | *Aldh1a2* | 7.67E-03 |
| 19 | *Marf1* | 1.70E-23 | 69 | *Ythdf2* | 4.65E-07 | 119 | *Lgr4* | 6.47E-04 | 169 | *Ccnd3* | 7.79E-03 |
| 20 | *Rpgrip1* | 6.95E-23 | 70 | *Ubn2* | 6.22E-07 | 120 | *Elavl2* | 6.64E-04 | 170 | *Slit2* | 8.36E-03 |
| 21 | *Rrn3* | 8.99E-23 | 71 | *Znrf3* | 6.66E-07 | 121 | *Kitl* | 6.65E-04 | 171 | *Golga7b* | 8.55E-03 |
| 22 | *Spry2* | 7.95E-22 | 72 | *Mn1* | 9.62E-07 | 122 | *Sept2* | 6.83E-04 | 172 | *Ank3* | 9.34E-03 |
| 23 | *Insig1* | 2.86E-19 | 73 | *Lsm14b* | 1.15E-06 | 123 | *Mtmr2* | 8.65E-04 | 173 | *Mast4* | 9.77E-03 |
| 24 | *Mcmbp* | 5.29E-19 | 74 | *Runx1t1* | 1.15E-06 | 124 | *Efna5* | 8.76E-04 | 174 | *Gm2694* | 1.00E-02 |
| 25 | *Pdcd4* | 1.37E-18 | 75 | *Supt16* | 1.25E-06 | 125 | *Jund* | 9.20E-04 | | | |

| | | | | | | | | |
|---|---|---|---|---|---|---|---|---|
| 26 | *Rbm15* | 2.91E-18 | 76 | *Picalm* | 1.43E-06 | 126 | *Csnk1e* | 1.04E-03 |
| 27 | *Gatad1* | 2.95E-18 | 77 | *Cux2* | 1.47E-06 | 127 | *Ado* | 1.18E-03 |
| 28 | *Wtap* | 3.51E-18 | 78 | *Zfp207* | 1.65E-06 | 128 | *Mex3a* | 1.23E-03 |
| 29 | *Dctn4* | 6.88E-18 | 79 | *Ptp4a2* | 3.09E-06 | 129 | *Acsl4* | 1.25E-03 |
| 30 | *Micu3* | 9.81E-18 | 80 | *Cdv3* | 3.11E-06 | 130 | *Mau2* | 1.53E-03 |
| 31 | *Sec23ip* | 5.24E-17 | 81 | *Yme1l1* | 7.00E-06 | 131 | *Ptma* | 1.80E-03 |
| 32 | *Yod1* | 7.72E-17 | 82 | *Atxn1* | 8.03E-06 | 132 | *Brd8* | 1.80E-03 |
| 33 | *Birc6* | 4.44E-16 | 83 | *Dusp3* | 1.01E-05 | 133 | *Cbll1* | 1.89E-03 |
| 34 | *Ppm1h* | 6.57E-16 | 84 | *Zbtb21* | 1.17E-05 | 134 | *Enah* | 2.02E-03 |
| 35 | *Ncan* | 1.39E-15 | 85 | *Nos1* | 1.21E-05 | 135 | *Zeb2* | 2.17E-03 |
| 36 | *Set* | 3.09E-15 | 86 | *Kcnip4* | 1.25E-05 | 136 | *Srpk1* | 2.20E-03 |
| 37 | *Ncs1* | 3.99E-15 | 87 | *Sirt6* | 1.28E-05 | 137 | *Grem2* | 2.40E-03 |
| 38 | *Skil* | 2.66E-14 | 88 | *Rap2b* | 1.42E-05 | 138 | *Tspyl5* | 2.41E-03 |
| 39 | *Nras* | 3.78E-14 | 89 | *Usp22* | 1.67E-05 | 139 | *Vstm2l* | 2.41E-03 |
| 40 | *Ric1* | 3.36E-12 | 90 | *Dfna5* | 1.69E-05 | 140 | *Med1* | 2.41E-03 |
| 41 | *Abhd2* | 6.91E-12 | 91 | *Mtf2* | 2.14E-05 | 141 | *Slitrk2* | 2.42E-03 |
| 42 | *Pan3* | 2.75E-11 | 92 | *Entpd7* | 3.69E-05 | 142 | *Zfhx3* | 2.60E-03 |
| 43 | *Slc39a1* | 3.07E-11 | 93 | *Hoxd10* | 3.79E-05 | 143 | *Med13l* | 3.19E-03 |
| 44 | *Clasp2* | 3.41E-11 | 94 | *Ppp1r16b* | 4.37E-05 | 144 | *Ccdc6* | 3.20E-03 |
| 45 | *Klf13* | 3.69E-11 | 95 | *Tbck* | 4.37E-05 | 145 | *Gnl3l* | 3.38E-03 |
| 46 | *Tmf1* | 1.77E-10 | 96 | *Dpf1* | 4.37E-05 | 146 | *Fam8a1* | 3.57E-03 |
| 47 | *Zscan26* | 2.00E-10 | 97 | *Hmgcr* | 5.45E-05 | 147 | *Stmn2* | 4.15E-03 |
| 48 | *Lzts1* | 2.16E-10 | 98 | *Casz1* | 6.57E-05 | 148 | *Smg7* | 4.68E-03 |
| 49 | *Amt* | 2.34E-10 | 99 | *Whsc1l1* | 1.01E-04 | 149 | *Prrc2c* | 4.79E-03 |
| 50 | *Kdelr2* | 4.00E-10 | 100 | *Nmt2* | 1.06E-04 | 150 | *Hoxa10* | 4.79E-03 |

**Supplementary Table 5.** Top 20 significant GO Biological Process terms using *Kdm6b* specific SGS-responsive genes from WT motor neurons in the embryonic mouse spine dataset.

| **Term** | **Overlap** | **P-value** | **Adj. P-value** | **Genes** |
|---|---|---|---|---|
| *Positive Regulation Of Transcription By RNA Polymerase II (GO:0045944)* | 23/938 | 7.82E-06 | 5.19E-03 | *KDM6B;MED1;KMT2D;ACTR2;GTF2A1;CASZ1;ZFHX3; JUND;TMF1;NCOA3;SOX11;NR2F2;FOXO3;ETV4; HOXC10;HOXA10;ZEB2;NFIA;HSF2;NOS1;BRD8;PTMA;SOX4* |
| *Regulation Of DNA-templated Transcription (GO:0006355)* | 36/1922 | 8.83E-06 | 5.19E-03 | *KMT2D;KHDRBS1;CASZ1;SET;ZBTB21;FOXO3;HOXD10; ACTB;HOXC10;ELAVL2;HOXA10;ATXN1;HSF2;DPF1;NOS1; BRD8;SOX4;RUNX1T1;MED1;ZFHX3;JUND;RBM15;KLF13; CBX3;BCL11A;USP22;F2R;NR2F2;ETV4; ZEB2;NFIA;MTF2; PDCD4;ZSCAN26;SP8;PICALM* |
| *Positive Regulation Of DNA-templated Transcription (GO:0045893)* | 27/1243 | 1.10E-05 | 5.19E-03 | *KMT2D;GTF2A1;CASZ1;TMF1;FOXO3;ACTB;HOXC10; HOXA10;HSF2;NOS1;BRD8;SOX4;KDM6B;MED1;ACTR2; ZFHX3;JUND;NCOA3;USP22;F2R;SOX11; NR2F2;ETV4; ZEB2;NFIA;PTMA;PICALM* |
| *Regulation Of Microtubule Cytoskeleton Organization (GO:0070507)* | 5/40 | 2.49E-05 | 6.73E-03 | *GSK3B;STMN2;STMN3;EFNA5;CLASP2* |
| *Positive Regulation Of Insulin Secretion (GO:0032024)* | 5/40 | 2.49E-05 | 6.73E-03 | *LRRC8A;ACSL4;SIRT6;SOX4;MCU* |
| *Regulation Of Transcription By RNA Polymerase II (GO:0006357)* | 36/2028 | 2.84E-05 | 6.73E-03 | *KMT2D;GTF2A1;CASZ1;TMF1;ZBTB21;FOXO3;HOXD10; ACTB;HOXC10;HOXA10;ATXN1;CUX2;HSF2;DPF1;NOS1; BRD8;SOX4;KDM6B;MED1;ACTR2;ZFHX3;JUND;KLF13; CBX3;BCL11A;NCOA3;USP22;SOX11; SIRT6;NR2F2; ETV4;ZEB2;NFIA;ZSCAN26;SP8;PTMA* |
| *Positive Regulation Of Peptide Hormone Secretion (GO:0090277)* | 5/46 | 4.97E-05 | 1.01E-02 | *LRRC8A;ACSL4;SIRT6;SOX4;MCU* |
| *Regulation Of Insulin Secretion (GO:0050796)* | 6/87 | 1.16E-04 | 2.06E-02 | *ACSL4;SIRT6;LRRC8A;EFNA5;SOX4;MCU* |
| *Positive Regulation Of Keratinocyte Differentiation (GO:0045618)* | 3/13 | 1.77E-04 | 2.79E-02 | *MED1;NCOA3;ETV4* |
| *Regulation Of Dephosphorylation (GO:0035303)* | 3/15 | 2.77E-04 | 3.94E-02 | *PPP1R16B;MTMR2;SMG7* |
| *Positive Regulation Of Epidermal Cell Differentiation (GO:0045606)* | 3/18 | 4.88E-04 | 6.30E-02 | *MED1;NCOA3;ETV4* |

| | | | | |
|---|---|---|---|---|
| *Positive Regulation Of Protein Secretion (GO:0050714)* | 5/76 | 5.42E-04 | 6.42E-02 | *ACSL4;SIRT6;LRRC8A;SOX4;MCU* |
| *Regulation Of Neuron Differentiation (GO:0045664)* | 5/79 | 6.47E-04 | 6.52E-02 | *MED1;CASZ1;ZFHX3;NTF3;SOX11* |
| *Striated Muscle Cell Differentiation (GO:0051146)* | 3/20 | 6.73E-04 | 6.52E-02 | *KDM6B;MYEF2;SIRT6* |
| *Regulation Of Focal Adhesion Disassembly (GO:0120182)* | 2/5 | 7.48E-04 | 6.52E-02 | *DUSP3;IQSEC1* |
| *Positive Regulation Of Focal Adhesion Disassembly (GO:0120183)* | 2/5 | 7.48E-04 | 6.52E-02 | *DUSP3;IQSEC1* |
| *Mitochondrial Calcium Ion Homeostasis (GO:0051560)* | 3/21 | 7.80E-04 | 6.52E-02 | *MICU3;PDZD8;MCU* |
| *Regulation Of Double-Strand Break Repair (GO:2000779)* | 5/88 | 1.06E-03 | 8.34E-02 | *ACTR2;DPF1;SIRT6;BRD8;OTUB1* |
| *Noradrenergic Neuron Differentiation (GO:0003357)* | 2/6 | 1.12E-03 | 8.35E-02 | *SOX11;SOX4* |
| *Regulation Of Alternative mRNA Splicing, Via Spliceosome (GO:0000381)* | 4/53 | 1.19E-03 | 8.44E-02 | *KHDRBS1;RBFOX1;RBM15;WTAP* |

**Supplementary Table 6.** *IL7R* specific SGS-responsive genes from CD4+ T cells in the PBMC20K dataset.

| Rank | Gene | FDR | Rank | Gene | FDR | Rank | Gene | FDR | Rank | Gene | FDR |
|---|---|---|---|---|---|---|---|---|---|---|---|
| 1 | *IL7R* | 0.00E+00 | 51 | *AC005224.2* | 5.11E-05 | 101 | *TNFRSF1B* | 1.58E-03 | 151 | *PARP14* | 6.71E-03 |
| 2 | *FOXP3* | 8.24E-57 | 52 | *OXNAD1* | 5.11E-05 | 102 | *ID2* | 1.58E-03 | 152 | *PRKCQ-AS1* | 6.91E-03 |
| 3 | *IKZF2* | 1.65E-45 | 53 | *NPM1* | 6.12E-05 | 103 | *VPS26C* | 1.58E-03 | 153 | *MAML2* | 6.92E-03 |
| 4 | *LINC02694* | 4.64E-41 | 54 | *RBMS1* | 8.78E-05 | 104 | *ACSL6* | 1.62E-03 | 154 | *TMEM71* | 6.92E-03 |
| 5 | *ANXA1* | 1.03E-19 | 55 | *GIMAP4* | 9.59E-05 | 105 | *TIGIT* | 1.63E-03 | 155 | *LST1* | 7.09E-03 |
| 6 | *RTKN2* | 3.51E-16 | 56 | *CTSW* | 1.11E-04 | 106 | *LINC00623* | 1.68E-03 | 156 | *ARHGAP5* | 7.24E-03 |
| 7 | *CTLA4* | 4.50E-15 | 57 | *UQCRB* | 1.11E-04 | 107 | *RGS19* | 1.81E-03 | 157 | *TRADD* | 7.47E-03 |
| 8 | *IL2RA* | 8.30E-14 | 58 | *ARPC1B* | 1.45E-04 | 108 | *AKT3* | 1.92E-03 | 158 | *PRMT2* | 7.47E-03 |
| 9 | *CCR4* | 1.34E-13 | 59 | *SOS1* | 1.45E-04 | 109 | *IFI44* | 2.05E-03 | 159 | *TRABD2A* | 7.91E-03 |
| 10 | *CYTOR* | 2.49E-11 | 60 | *TOMM7* | 1.57E-04 | 110 | *EIF3E* | 2.07E-03 | 160 | *JUNB* | 8.46E-03 |
| 11 | *HLA-DRB1* | 3.41E-11 | 61 | *CD226* | 1.57E-04 | 111 | *BCAS2* | 2.09E-03 | 161 | *APPL1* | 8.46E-03 |
| 12 | *HPGD* | 2.07E-10 | 62 | *ZC2HC1A* | 1.57E-04 | 112 | *GPSM3* | 2.16E-03 | 162 | *CD74* | 8.66E-03 |
| 13 | *CD48* | 5.84E-09 | 63 | *PIK3IP1* | 1.57E-04 | 113 | *SSR3* | 2.16E-03 | 163 | *KLRB1* | 8.97E-03 |
| 14 | *ITM2B* | 4.97E-08 | 64 | *CCSER2* | 1.63E-04 | 114 | *LYSMD2* | 2.22E-03 | 164 | *NIBAN1* | 8.97E-03 |
| 15 | *PDCD4* | 8.50E-08 | 65 | *TESPA1* | 1.63E-04 | 115 | *TTN* | 2.24E-03 | 165 | *SOCS3* | 8.98E-03 |
| 16 | *MDFIC* | 1.01E-07 | 66 | *PLAAT4* | 1.68E-04 | 116 | *COX4I1* | 2.24E-03 | 166 | *LRRN3* | 9.43E-03 |
| 17 | *NOSIP* | 1.06E-07 | 67 | *COQ10B* | 1.96E-04 | 117 | *GPR171* | 2.24E-03 | 167 | *CNOT2* | 9.82E-03 |
| 18 | *HINT1* | 1.33E-07 | 68 | *SNHG29* | 1.96E-04 | 118 | *LGALS3* | 2.25E-03 | 168 | *LMO4* | 9.84E-03 |
| 19 | *TCF7* | 2.86E-07 | 69 | *PLAC8* | 2.02E-04 | 119 | *NCOA7* | 2.45E-03 | 169 | *HIPK2* | 9.84E-03 |
| 20 | *HLA-DPB1* | 3.37E-07 | 70 | *LINC00426* | 2.19E-04 | 120 | *PLP2* | 2.53E-03 | | | |
| 21 | *GIMAP7* | 4.16E-07 | 71 | *CDKN2AIP* | 2.61E-04 | 121 | *SNHG7* | 2.78E-03 | | | |
| 22 | *JPT1* | 4.99E-07 | 72 | *CSNK1G2* | 2.62E-04 | 122 | *FYB1* | 2.83E-03 | | | |
| 23 | *TRAT1* | 1.46E-06 | 73 | *DOK2* | 2.74E-04 | 123 | *PIM1* | 2.83E-03 | | | |
| 24 | *CCR10* | 1.51E-06 | 74 | *CD79B* | 3.11E-04 | 124 | *ACTR3* | 2.83E-03 | | | |
| 25 | *GAS5* | 1.98E-06 | 75 | *DYNLL1* | 3.21E-04 | 125 | *PCED1B-AS1* | 2.87E-03 | | | |

| | | | | | | | | |
|---|---|---|---|---|---|---|---|---|
| 26 | *HLA-DPA1* | 2.23E-06 | 76 | *CAMK4* | 3.42E-04 | 126 | *RNASEH2B* | 2.98E-03 |
| 27 | *ZFP36L2* | 2.91E-06 | 77 | *AP3M2* | 3.49E-04 | 127 | *HLA-DRA* | 3.18E-03 |
| 28 | *GIMAP8* | 3.70E-06 | 78 | *MIAT* | 3.50E-04 | 128 | *TAGAP* | 3.18E-03 |
| 29 | *MYBL1* | 7.24E-06 | 79 | *SATB1* | 4.77E-04 | 129 | *SORL1* | 3.31E-03 |
| 30 | *TNFAIP8* | 7.53E-06 | 80 | *LAIR2* | 4.83E-04 | 130 | *MID1IP1* | 3.31E-03 |
| 31 | *RIPOR2* | 7.73E-06 | 81 | *PNRC1* | 4.89E-04 | 131 | *MCUB* | 3.51E-03 |
| 32 | *CORO1A* | 9.36E-06 | 82 | *CDK6* | 5.05E-04 | 132 | *SEMA4D* | 3.57E-03 |
| 33 | *TIMP1* | 1.08E-05 | 83 | *GIMAP1* | 5.05E-04 | 133 | *IFITM2* | 3.60E-03 |
| 34 | *FXYD5* | 1.53E-05 | 84 | *TRIM44* | 5.47E-04 | 134 | *NT5C* | 3.61E-03 |
| 35 | *INPP4B* | 1.72E-05 | 85 | *MAL* | 7.02E-04 | 135 | *CST7* | 3.70E-03 |
| 36 | *MAP3K1* | 1.72E-05 | 86 | *TC2N* | 7.49E-04 | 136 | *CANX* | 3.90E-03 |
| 37 | *LIMA1* | 1.93E-05 | 87 | *PASK* | 8.15E-04 | 137 | *PRKD3* | 3.90E-03 |
| 38 | *EEF2* | 1.94E-05 | 88 | *CD69* | 8.23E-04 | 138 | *CHD7* | 4.10E-03 |
| 39 | *IL6ST* | 2.14E-05 | 89 | *CD40LG* | 8.32E-04 | 139 | *PARP8* | 4.43E-03 |
| 40 | *HOPX* | 2.25E-05 | 90 | *CRBN* | 8.86E-04 | 140 | *KLRG1* | 4.57E-03 |
| 41 | *SERINC5* | 2.29E-05 | 91 | *UBQLN2* | 9.05E-04 | 141 | *CD96* | 4.76E-03 |
| 42 | *LPIN2* | 2.31E-05 | 92 | *KLF2* | 1.02E-03 | 142 | *C12orf75* | 4.82E-03 |
| 43 | *NFKBIA* | 2.37E-05 | 93 | *ERN1* | 1.02E-03 | 143 | *OAS1* | 4.88E-03 |
| 44 | *FKBP5* | 2.37E-05 | 94 | *SERP1* | 1.10E-03 | 144 | *XBP1* | 4.88E-03 |
| 45 | *ANK3* | 2.41E-05 | 95 | *TGFBR2* | 1.10E-03 | 145 | *NCBP2AS2* | 4.88E-03 |
| 46 | *PTGER2* | 2.65E-05 | 96 | *NOP53* | 1.21E-03 | 146 | *EAPP* | 4.89E-03 |
| 47 | *SSR2* | 3.79E-05 | 97 | *SCML4* | 1.24E-03 | 147 | *AC004556.3* | 5.56E-03 |
| 48 | *GIMAP5* | 3.79E-05 | 98 | *CCL5* | 1.27E-03 | 148 | *DPP4* | 5.60E-03 |
| 49 | *SLC40A1* | 3.93E-05 | 99 | *PABPC1* | 1.32E-03 | 149 | *ZFAS1* | 5.71E-03 |
| 50 | *SNHG6* | 4.34E-05 | 100 | *CDC14A* | 1.45E-03 | 150 | *IFI44L* | 6.71E-03 |

**Supplementary Table 7.** Significant GO Biological Process Terms using the top 100 *IL7R* specific SGS-responsive genes from CD4+ T cells in the PBMC20K dataset.

| **Term** | **Overlap** | **P-value** | **Adj. P-value** | **Genes** |
|---|---|---|---|---|
| *Regulation Of T Cell Proliferation (GO:0042129)* | 7/77 | 1.14E-07 | 1.22E-04 | *ANXA1;CD40LG;CCL5;IL6ST;FOXP3;HLA-DRB1;HLA-DPA1* |
| *Regulation Of T Cell Tolerance Induction (GO:0002664)* | 3/6 | 2.40E-06 | 1.29E-03 | *IL2RA;FOXP3;TGFBR2* |
| *Positive Regulation Of T Cell Activation (GO:0050870)* | 6/107 | 1.61E-05 | 4.93E-03 | *ANXA1;CD40LG;CCL5;IL6ST;HLA-DRB1;HLA-DPA1* |
| *Positive Regulation Of T Cell Proliferation (GO:0042102)* | 5/65 | 1.84E-05 | 4.93E-03 | *ANXA1;CD40LG;CCL5;IL6ST;HLA-DPA1* |
| *Cellular Response To Cytokine Stimulus (GO:0071345)* | 9/308 | 2.42E-05 | 5.18E-03 | *NFKBIA;RIPOR2;CCL5;TCF7;IL6ST;IL7R;SOS1; ZFP36L2;HLA-DPA1* |
| *Positive Regulation Of T Cell Receptor Signaling Pathway (GO:0050862)* | 3/13 | 3.34E-05 | 5.30E-03 | *TESPA1;TRAT1;CD226* |
| *Positive Regulation Of Lymphocyte Proliferation (GO:0050671)* | 5/74 | 3.46E-05 | 5.30E-03 | *ANXA1;CD40LG;CCL5;IL6ST;HLA-DPA1* |
| *Positive Regulation Of Signal Transduction (GO:0009967)* | 8/266 | 5.71E-05 | 7.65E-03 | *ERN1;CRBN;CD226;IL6ST;IL7R;CDKN2AIP;TRIM44;KLF2* |
| *T Cell Differentiation (GO:0030217)* | 4/46 | 8.17E-05 | 9.73E-03 | *ANXA1;TCF7;FOXP3;ZFP36L2* |
| *Positive Regulation Of Antigen Receptor-Mediated Signaling Pathway (GO:0050857)* | 3/20 | 1.30E-04 | 1.39E-02 | *TESPA1;TRAT1;CD226* |
| *Regulation Of Interleukin-4 Production (GO:0032673)* | 3/23 | 2.00E-04 | 1.87E-02 | *CD40LG;FOXP3;HLA-DRB1* |
| *Positive Regulation Of Translation (GO:0045727)* | 5/108 | 2.10E-04 | 1.87E-02 | *NPM1;CCL5;PABPC1;PASK;EEF2* |
| *Negative Regulation Of Cellular Process (GO:0048523)* | 10/537 | 3.56E-04 | 2.94E-02 | *RIPOR2;NPM1;CDK6;HPGD;DYNLL1;FOXP3;MYBL1; PLAAT4;CDKN2AIP;HOPX* |
| *Regulation Of T Cell Receptor Signaling Pathway (GO:0050856)* | 3/34 | 6.48E-04 | 4.97E-02 | *TESPA1;TRAT1;CD226* |
| *Positive Regulation Of Immune Response (GO:0050778)* | 4/80 | 6.95E-04 | 4.97E-02 | *CCL5;CD226;IL6ST;HLA-DRB1* |

**Supplementary Table 8.** Top 200 *STAT1* specific SGS-responsive genes from CD4+ T cells in the PBMC5K dataset.

| Rank | Gene | FDR | Rank | Gene | FDR | Rank | Gene | FDR | Rank | Gene | FDR |
|---|---|---|---|---|---|---|---|---|---|---|---|
| 1 | *STAT1* | 0.00E+00 | 51 | *SEC13* | 6.89E-04 | 101 | *BRAF* | 2.53E-03 | 151 | *SNRNP40* | 3.98E-03 |
| 2 | *FAU* | 2.97E-08 | 52 | *SLC38A1* | 7.00E-04 | 102 | *GNAI3* | 2.58E-03 | 152 | *CDYL* | 4.01E-03 |
| 3 | *GBP5* | 4.85E-08 | 53 | *HECTD4* | 7.00E-04 | 103 | *ARHGAP25* | 2.62E-03 | 153 | *TBC1D14* | 4.14E-03 |
| 4 | *EPSTI1* | 4.85E-08 | 54 | *MAP3K1* | 7.11E-04 | 104 | *FAM214A* | 2.62E-03 | 154 | *CAAP1* | 4.14E-03 |
| 5 | *NOP53* | 8.25E-08 | 55 | *MAP3K4* | 7.64E-04 | 105 | *DDHD1* | 2.64E-03 | 155 | *ZFP91* | 4.14E-03 |
| 6 | *ARHGAP15* | 3.20E-07 | 56 | *TMEM161B* | 7.71E-04 | 106 | *APOL3* | 2.64E-03 | 156 | *GPCPD1* | 4.14E-03 |
| 7 | *PARP9* | 3.20E-07 | 57 | *PTBP2* | 7.90E-04 | 107 | *PPP4R3B* | 2.72E-03 | 157 | *KYAT3* | 4.16E-03 |
| 8 | *RACK1* | 3.25E-07 | 58 | *MAN2A1* | 7.90E-04 | 108 | *CAMK4* | 2.72E-03 | 158 | *EIF1* | 4.26E-03 |
| 9 | *EEF1A1* | 6.95E-07 | 59 | *CD226* | 9.09E-04 | 109 | *CAPZA2* | 2.82E-03 | 159 | *LATS1* | 4.32E-03 |
| 10 | *SP100* | 1.67E-06 | 60 | *DNAJC1* | 9.22E-04 | 110 | *RBM33* | 2.82E-03 | 160 | *DYNC1LI2* | 4.36E-03 |
| 11 | *GBP2* | 2.84E-06 | 61 | *GPRIN3* | 9.22E-04 | 111 | *ACTN4* | 2.92E-03 | 161 | *HEXA* | 4.37E-03 |
| 12 | *EEF1D* | 3.01E-06 | 62 | *PICALM* | 9.22E-04 | 112 | *PSME4* | 2.94E-03 | 162 | *DISC1* | 4.55E-03 |
| 13 | *UBA52* | 3.04E-06 | 63 | *CDC14A* | 9.40E-04 | 113 | *NCOA2* | 2.98E-03 | 163 | *PRKDC* | 4.55E-03 |
| 14 | *EEF1B2* | 3.16E-06 | 64 | *RNF185* | 9.46E-04 | 114 | *PTPRC* | 2.99E-03 | 164 | *PHF21A* | 4.55E-03 |
| 15 | *NLRC5* | 3.40E-06 | 65 | *MAPK8* | 1.03E-03 | 115 | *JADE2* | 2.99E-03 | 165 | *SNIP1* | 4.67E-03 |
| 16 | *SNHG29* | 5.83E-06 | 66 | *BTF3* | 1.11E-03 | 116 | *PARP12* | 3.01E-03 | 166 | *NIBAN1* | 4.67E-03 |
| 17 | *NACA* | 9.48E-06 | 67 | *STAU1* | 1.11E-03 | 117 | *NCSTN* | 3.17E-03 | 167 | *SLC4A7* | 4.67E-03 |
| 18 | *GBP4* | 1.90E-05 | 68 | *IFI16* | 1.12E-03 | 118 | *CYP20A1* | 3.17E-03 | 168 | *ELL2* | 4.67E-03 |
| 19 | *TPT1* | 2.57E-05 | 69 | *AC068587.4* | 1.14E-03 | 119 | *XRN1* | 3.17E-03 | 169 | *ATXN1* | 4.67E-03 |
| 20 | *PTMA* | 2.99E-05 | 70 | *CHST11* | 1.14E-03 | 120 | *TRIM23* | 3.17E-03 | 170 | *VWA8* | 4.67E-03 |
| 21 | *TOMM7* | 3.50E-05 | 71 | *CEMIP2* | 1.24E-03 | 121 | *ATP6V1H* | 3.17E-03 | 171 | *ABHD13* | 4.68E-03 |
| 22 | *GAS5* | 3.57E-05 | 72 | *DIS3* | 1.24E-03 | 122 | *CTSH* | 3.17E-03 | 172 | *MED13* | 4.68E-03 |
| 23 | *MEF2A* | 3.57E-05 | 73 | *PPP1R16B* | 1.24E-03 | 123 | *XAF1* | 3.17E-03 | 173 | *APOL6* | 4.68E-03 |
| 24 | *DPYD* | 4.34E-05 | 74 | *TXNIP* | 1.27E-03 | 124 | *MECP2* | 3.23E-03 | 174 | *ZFC3H1* | 4.74E-03 |
| 25 | *SPPL2A* | 4.45E-05 | 75 | *UTRN* | 1.27E-03 | 125 | *WBP11* | 3.26E-03 | 175 | *ATP2B1* | 4.74E-03 |

| | | | | | | | | | | | |
|---|---|---|---|---|---|---|---|---|---|---|---|
| 26 | *DNAJC3* | 4.45E-05 | 76 | *WDR48* | 1.46E-03 | 126 | *WDFY1* | 3.32E-03 | 176 | *HDAC5* | 4.78E-03 |
| 27 | *RAB8B* | 5.40E-05 | 77 | *ARNTL* | 1.47E-03 | 127 | *CDV3* | 3.32E-03 | 177 | *ELMO1* | 4.79E-03 |
| 28 | *YTHDF3* | 6.41E-05 | 78 | *RAPGEF1* | 1.49E-03 | 128 | *ACAP2* | 3.32E-03 | 178 | *ANKRD28* | 4.84E-03 |
| 29 | *AKT3* | 8.86E-05 | 79 | *UQCRC2* | 1.54E-03 | 129 | *COMMD6* | 3.32E-03 | 179 | *OTULINL* | 4.87E-03 |
| 30 | *BTG1* | 1.17E-04 | 80 | *NDUFA4* | 1.56E-03 | 130 | *RASA3* | 3.32E-03 | 180 | *SLCO3A1* | 4.97E-03 |
| 31 | *ULK4* | 1.46E-04 | 81 | *MBD5* | 1.63E-03 | 131 | *TRPM7* | 3.32E-03 | 181 | *FAM91A1* | 5.03E-03 |
| 32 | *CDC42SE2* | 1.46E-04 | 82 | *SKAP1* | 1.63E-03 | 132 | *CCDC84* | 3.39E-03 | 182 | *USP48* | 5.24E-03 |
| 33 | *FRS2* | 1.46E-04 | 83 | *MOB1A* | 1.77E-03 | 133 | *ERP44* | 3.58E-03 | 183 | *SENP7* | 5.25E-03 |
| 34 | *LPP* | 1.85E-04 | 84 | *MLLT10* | 1.77E-03 | 134 | *C5orf56* | 3.58E-03 | 184 | *HIP1R* | 5.25E-03 |
| 35 | *TUBA1B* | 1.97E-04 | 85 | *CDC123* | 1.86E-03 | 135 | *SHPRH* | 3.58E-03 | 185 | *TET2* | 5.29E-03 |
| 36 | *SYNE2* | 2.64E-04 | 86 | *COPS2* | 1.86E-03 | 136 | *THUMPD2* | 3.64E-03 | 186 | *STAT2* | 5.29E-03 |
| 37 | *SUCLA2* | 3.00E-04 | 87 | *ANKRD44* | 1.86E-03 | 137 | *BMS1* | 3.64E-03 | 187 | *LRRC37B* | 5.44E-03 |
| 38 | *EIF3K* | 3.00E-04 | 88 | *ZEB1* | 1.86E-03 | 138 | *IGF2R* | 3.79E-03 | 188 | *SERTAD2* | 5.59E-03 |
| 39 | *TRIM38* | 3.08E-04 | 89 | *ABCC1* | 1.86E-03 | 139 | *FAR1* | 3.79E-03 | 189 | *SPEF2* | 5.60E-03 |
| 40 | *USP15* | 3.34E-04 | 90 | *GTF2I* | 1.93E-03 | 140 | *ARHGAP17* | 3.79E-03 | 190 | *ECD* | 5.65E-03 |
| 41 | *EEF1G* | 3.55E-04 | 91 | *TXNDC11* | 1.93E-03 | 141 | *RAD54L2* | 3.79E-03 | 191 | *DLG1* | 5.66E-03 |
| 42 | *SREK1* | 3.56E-04 | 92 | *EEF2* | 1.93E-03 | 142 | *DOCK10* | 3.88E-03 | 192 | *PSMD12* | 5.69E-03 |
| 43 | *PFDN5* | 3.85E-04 | 93 | *HIST1H1E* | 2.02E-03 | 143 | *PLCL2* | 3.90E-03 | 193 | *OSBPL9* | 5.73E-03 |
| 44 | *ZFAND3* | 3.97E-04 | 94 | *CAPN2* | 2.05E-03 | 144 | *PIGG* | 3.90E-03 | 194 | *SNHG5* | 5.73E-03 |
| 45 | *SPIDR* | 3.97E-04 | 95 | *SOS1* | 2.05E-03 | 145 | *SNHG32* | 3.90E-03 | 195 | *HELB* | 5.73E-03 |
| 46 | *PCNX1* | 4.20E-04 | 96 | *EXOC2* | 2.05E-03 | 146 | *CGAS* | 3.90E-03 | 196 | *MIF* | 5.73E-03 |
| 47 | *PDIK1L* | 4.68E-04 | 97 | *PREX1* | 2.05E-03 | 147 | *NSMCE4A* | 3.90E-03 | 197 | *EIF2AK2* | 5.76E-03 |
| 48 | *RNPC3* | 5.64E-04 | 98 | *ADD1* | 2.15E-03 | 148 | *RNF10* | 3.90E-03 | 198 | *HLTF* | 5.79E-03 |
| 49 | *FTL* | 5.78E-04 | 99 | *CNOT6L* | 2.15E-03 | 149 | *AP3D1* | 3.90E-03 | 199 | *AGTPBP1* | 5.79E-03 |
| 50 | *SIN3B* | 6.33E-04 | 100 | *TUT7* | 2.46E-03 | 150 | *PBX4* | 3.95E-03 | 200 | *ATP6V0D1* | 5.79E-03 |

**Supplementary Table 9.** Top 200 *STAT1* specific SGS-responsive genes from CD4+ T cells in the PBMC10K dataset.

| Rank | Gene | FDR | Rank | Gene | FDR | Rank | Gene | FDR | Rank | Gene | FDR |
|---|---|---|---|---|---|---|---|---|---|---|---|
| 1 | *STAT1* | 0.00E+00 | 51 | *ANXA1* | 1.50E-05 | 101 | *EMP3* | 3.35E-04 | 151 | *TMEM179B* | 1.47E-03 |
| 2 | *GBP1* | 6.53E-23 | 52 | *DDX58* | 1.84E-05 | 102 | *PSMA3* | 3.58E-04 | 152 | *TPGS2* | 1.47E-03 |
| 3 | *PSMB9* | 1.74E-13 | 53 | *YWHAZ* | 1.87E-05 | 103 | *ACTN4* | 3.61E-04 | 153 | *OGFR* | 1.48E-03 |
| 4 | *SYNE2* | 1.74E-13 | 54 | *BTG1* | 2.03E-05 | 104 | *SETX* | 3.77E-04 | 154 | *USP48* | 1.50E-03 |
| 5 | *IRF1* | 7.10E-13 | 55 | *TYMP* | 2.05E-05 | 105 | *IFI44* | 3.89E-04 | 155 | *CERS2* | 1.51E-03 |
| 6 | *PARP9* | 1.12E-12 | 56 | *LGALS1* | 3.54E-05 | 106 | *CHMP5* | 3.89E-04 | 156 | *CARD16* | 1.52E-03 |
| 7 | *LY6E* | 5.04E-12 | 57 | *STAT2* | 4.14E-05 | 107 | *TRIM21* | 3.89E-04 | 157 | *MRPS36* | 1.56E-03 |
| 8 | *XAF1* | 8.12E-12 | 58 | *NEAT1* | 4.47E-05 | 108 | *CPNE1* | 3.89E-04 | 158 | *NUTM2B-AS1* | 1.56E-03 |
| 9 | *TAP1* | 9.76E-12 | 59 | *MX2* | 4.47E-05 | 109 | *SP100* | 3.94E-04 | 159 | *EEF1G* | 1.59E-03 |
| 10 | *GBP2* | 1.32E-11 | 60 | *TRANK1* | 5.55E-05 | 110 | *EFTUD2* | 3.97E-04 | 160 | *SH3GLB1* | 1.61E-03 |
| 11 | *EPSTI1* | 1.32E-11 | 61 | *ADGRE5* | 5.81E-05 | 111 | *FAU* | 4.02E-04 | 161 | *PHF11* | 1.61E-03 |
| 12 | *PSME2* | 1.88E-11 | 62 | *S100A4* | 5.97E-05 | 112 | *C17orf49* | 4.24E-04 | 162 | *AFF1* | 1.62E-03 |
| 13 | *UBE2L6* | 5.38E-11 | 63 | *S100A10* | 6.03E-05 | 113 | *IRF9* | 4.42E-04 | 163 | *FOSB* | 1.67E-03 |
| 14 | *IFI44L* | 2.63E-10 | 64 | *EEF1D* | 6.75E-05 | 114 | *DPP4* | 4.90E-04 | 164 | *MAPKAPK5* | 1.72E-03 |
| 15 | *GBP4* | 2.63E-10 | 65 | *HOPX* | 7.49E-05 | 115 | *UBA52* | 4.90E-04 | 165 | *MYD88* | 1.79E-03 |
| 16 | *ISG15* | 7.59E-10 | 66 | *CLIC1* | 8.21E-05 | 116 | *PARP14* | 4.91E-04 | 166 | *GLUL* | 1.82E-03 |
| 17 | *SAMD9L* | 7.59E-10 | 67 | *HEATR1* | 8.34E-05 | 117 | *VIM* | 4.91E-04 | 167 | *PDCD5* | 1.82E-03 |
| 18 | *PSME1* | 1.01E-09 | 68 | *OTUD4* | 8.34E-05 | 118 | *GBP3* | 5.00E-04 | 168 | *SMDT1* | 1.82E-03 |
| 19 | *MYO1F* | 5.72E-09 | 69 | *SQOR* | 8.38E-05 | 119 | *PTGER4* | 5.14E-04 | 169 | *SLF2* | 1.86E-03 |
| 20 | *EIF2AK2* | 5.75E-09 | 70 | *PSENEN* | 9.30E-05 | 120 | *SHOC2* | 6.53E-04 | 170 | *EIF3J* | 1.86E-03 |
| 21 | *APOL6* | 9.17E-09 | 71 | *APOL3* | 1.01E-04 | 121 | *SAMHD1* | 6.54E-04 | 171 | *BTN3A2* | 1.90E-03 |
| 22 | *MT2A* | 1.33E-08 | 72 | *TRADD* | 1.02E-04 | 122 | *ERVK3-1* | 6.57E-04 | 172 | *TNIP2* | 1.90E-03 |
| 23 | *S100A11* | 1.66E-08 | 73 | *DTX3L* | 1.05E-04 | 123 | *LMO4* | 6.73E-04 | 173 | *EFR3A* | 1.99E-03 |
| 24 | *DDX60* | 2.45E-08 | 74 | *EEF1A1* | 1.09E-04 | 124 | *MOB1A* | 7.53E-04 | 174 | *MRPS22* | 1.99E-03 |
| 25 | *OAS1* | 2.45E-08 | 75 | *TRAFD1* | 1.11E-04 | 125 | *RACK1* | 7.53E-04 | 175 | *DOCK10* | 2.06E-03 |

| | | | | | | | | | | | |
|---|---|---|---|---|---|---|---|---|---|---|---|
| 26 | *GBP5* | 4.15E-08 | 76 | *GLS* | 1.12E-04 | 126 | *OAS2* | 7.80E-04 | 176 | *RAPGEF1* | 2.06E-03 |
| 27 | *RNF213* | 1.12E-07 | 77 | *SH3BP5* | 1.14E-04 | 127 | *AHNAK* | 8.27E-04 | 177 | *TRIM69* | 2.06E-03 |
| 28 | *NLRC5* | 1.15E-07 | 78 | *SAT1* | 1.20E-04 | 128 | *TPT1* | 8.38E-04 | 178 | *OPTN* | 2.13E-03 |
| 29 | *MX1* | 1.95E-07 | 79 | *ADAM19* | 1.29E-04 | 129 | *LAP3* | 9.17E-04 | 179 | *RIOK3* | 2.16E-03 |
| 30 | *CALM1* | 5.94E-07 | 80 | *PITPNB* | 1.29E-04 | 130 | *PHPT1* | 9.37E-04 | 180 | *RNF4* | 2.18E-03 |
| 31 | *HELZ2* | 1.31E-06 | 81 | *SPTAN1* | 1.35E-04 | 131 | *CYSLTR1* | 9.56E-04 | 181 | *SH3BGRL3* | 2.20E-03 |
| 32 | *CASP4* | 1.40E-06 | 82 | *FLNA* | 1.36E-04 | 132 | *IL10RA* | 9.68E-04 | 182 | *SNHG29* | 2.20E-03 |
| 33 | *IFI35* | 1.40E-06 | 83 | *AIDA* | 1.38E-04 | 133 | *NPDC1* | 9.91E-04 | 183 | *SLFN5* | 2.20E-03 |
| 34 | *CD74* | 1.95E-06 | 84 | *B2M* | 1.41E-04 | 134 | *VMP1* | 1.06E-03 | 184 | *SNUPN* | 2.27E-03 |
| 35 | *SARAF* | 1.96E-06 | 85 | *NBR1* | 1.41E-04 | 135 | *FUT7* | 1.07E-03 | 185 | *ADSS* | 2.28E-03 |
| 36 | *UBE2K* | 1.97E-06 | 86 | *TMSB10* | 1.44E-04 | 136 | *C5orf56* | 1.11E-03 | 186 | *EIF3A* | 2.36E-03 |
| 37 | *PSMB8* | 3.57E-06 | 87 | *AHR* | 1.71E-04 | 137 | *GATA3* | 1.11E-03 | 187 | *PSMD2* | 2.39E-03 |
| 38 | *TPM3* | 3.62E-06 | 88 | *CRYBG1* | 1.71E-04 | 138 | *DYNC1H1* | 1.15E-03 | 188 | *HERC5* | 2.39E-03 |
| 39 | *IL32* | 5.73E-06 | 89 | *SP140* | 1.86E-04 | 139 | *SPAG9* | 1.19E-03 | 189 | *NABP1* | 2.63E-03 |
| 40 | *CD3E* | 6.81E-06 | 90 | *SOS1* | 1.91E-04 | 140 | *GSTK1* | 1.20E-03 | 190 | *NDUFS1* | 2.69E-03 |
| 41 | *CRIP1* | 8.70E-06 | 91 | *RCBTB2* | 1.91E-04 | 141 | *SMIM14* | 1.23E-03 | 191 | *CD7* | 2.78E-03 |
| 42 | *CD99* | 1.00E-05 | 92 | *TRIM22* | 2.02E-04 | 142 | *DBNL* | 1.23E-03 | 192 | *ANXA5* | 2.78E-03 |
| 43 | *IFIH1* | 1.02E-05 | 93 | *FIBP* | 2.39E-04 | 143 | *PRKCQ* | 1.23E-03 | 193 | *ILF3* | 2.82E-03 |
| 44 | *PRDM1* | 1.11E-05 | 94 | *CXCR3* | 2.52E-04 | 144 | *IRF7* | 1.23E-03 | 194 | *OXSR1* | 2.84E-03 |
| 45 | *EEF1B2* | 1.13E-05 | 95 | *NIBAN1* | 2.64E-04 | 145 | *TOPBP1* | 1.28E-03 | 195 | *HLA-A* | 2.88E-03 |
| 46 | *TNFSF10* | 1.13E-05 | 96 | *NACA* | 2.83E-04 | 146 | *HERPUD1* | 1.28E-03 | 196 | *PRR14L* | 2.91E-03 |
| 47 | *ITGB1* | 1.13E-05 | 97 | *CHST12* | 3.23E-04 | 147 | *CTSS* | 1.35E-03 | 197 | *DESI2* | 2.95E-03 |
| 48 | *PARP11* | 1.13E-05 | 98 | *ANXA2* | 3.23E-04 | 148 | *GZMA* | 1.38E-03 | 198 | *NCL* | 2.95E-03 |
| 49 | *CAST* | 1.32E-05 | 99 | *CTSA* | 3.23E-04 | 149 | *ACTR3* | 1.44E-03 | 199 | *DARS* | 3.01E-03 |
| 50 | *CMTM6* | 1.40E-05 | 100 | *LCP1* | 3.30E-04 | 150 | *HSDL2* | 1.47E-03 | 200 | *HIST1H1E* | 3.08E-03 |

**Supplementary Table 10.** Top 200 *STAT1* specific SGS-responsive genes from CD4+ T cells in the PBMC20K dataset.

| Rank | Gene | FDR | Rank | Gene | FDR | Rank | Gene | FDR | Rank | Gene | FDR |
|---|---|---|---|---|---|---|---|---|---|---|---|
| 1 | *STAT1* | 0.00E+00 | 51 | *GZMA* | 1.18E-09 | 101 | *IFI16* | 1.58E-06 | 151 | *TRADD* | 1.71E-05 |
| 2 | *GBP1* | 4.11E-48 | 52 | *PIK3IP1* | 1.64E-09 | 102 | *SELL* | 1.60E-06 | 152 | *KDSR* | 1.81E-05 |
| 3 | *EPSTI1* | 2.52E-32 | 53 | *TMSB10* | 1.64E-09 | 103 | *MDFIC* | 1.72E-06 | 153 | *PLAAT4* | 1.96E-05 |
| 4 | *IRF1* | 8.56E-32 | 54 | *NIBAN1* | 2.11E-09 | 104 | *LGALS1* | 1.86E-06 | 154 | *SEC61B* | 2.09E-05 |
| 5 | *UBE2L6* | 7.85E-30 | 55 | *SP140* | 3.58E-09 | 105 | *TNFRSF4* | 1.89E-06 | 155 | *PTBP3* | 2.09E-05 |
| 6 | *PARP9* | 2.33E-29 | 56 | *SP100* | 3.80E-09 | 106 | *PRDM1* | 2.02E-06 | 156 | *LPGAT1* | 2.15E-05 |
| 7 | *PSMB9* | 1.45E-28 | 57 | *IFI6* | 4.52E-09 | 107 | *MAP4* | 2.12E-06 | 157 | *IL2RG* | 2.15E-05 |
| 8 | *GBP4* | 5.33E-26 | 58 | *PSMB8* | 5.39E-09 | 108 | *DDX60L* | 2.25E-06 | 158 | *CLIC1* | 2.41E-05 |
| 9 | *B2M* | 7.26E-25 | 59 | *FUT7* | 6.94E-09 | 109 | *PSMA6* | 2.85E-06 | 159 | *HMGN4* | 2.42E-05 |
| 10 | *GBP2* | 4.78E-24 | 60 | *ZCCHC2* | 7.02E-09 | 110 | *APOL2* | 2.90E-06 | 160 | *C1GALT1* | 2.44E-05 |
| 11 | *PSME2* | 5.66E-24 | 61 | *IFI44* | 8.36E-09 | 111 | *SETX* | 3.78E-06 | 161 | *C12orf75* | 2.44E-05 |
| 12 | *SAMD9L* | 1.10E-23 | 62 | *TYMP* | 1.17E-08 | 112 | *OAS2* | 4.33E-06 | 162 | *SLCO3A1* | 2.44E-05 |
| 13 | *NLRC5* | 3.21E-23 | 63 | *IRF9* | 1.55E-08 | 113 | *OPTN* | 4.67E-06 | 163 | *CD3G* | 2.49E-05 |
| 14 | *DTX3L* | 9.32E-21 | 64 | *GAS5* | 2.60E-08 | 114 | *HAPLN3* | 4.67E-06 | 164 | *CDC123* | 2.75E-05 |
| 15 | *IFI44L* | 1.09E-19 | 65 | *GLIPR1* | 3.24E-08 | 115 | *MAP3K1* | 5.17E-06 | 165 | *HOPX* | 2.89E-05 |
| 16 | *BTG1* | 2.12E-19 | 66 | *CALCOCO2* | 3.29E-08 | 116 | *LCP1* | 5.26E-06 | 166 | *SLC2A3* | 2.95E-05 |
| 17 | *TNFSF10* | 3.47E-19 | 67 | *RACK1* | 4.57E-08 | 117 | *CISH* | 5.36E-06 | 167 | *ERVK3-1* | 2.95E-05 |
| 18 | *GBP5* | 9.97E-19 | 68 | *HELZ2* | 5.89E-08 | 118 | *ARPC3* | 5.36E-06 | 168 | *IFITM1* | 3.47E-05 |
| 19 | *PARP14* | 1.53E-18 | 69 | *MYO1F* | 6.24E-08 | 119 | *PREX1* | 5.38E-06 | 169 | *PARP4* | 3.47E-05 |
| 20 | *PSME1* | 1.68E-18 | 70 | *OAS1* | 7.82E-08 | 120 | *USP46* | 5.62E-06 | 170 | *LAP3* | 3.54E-05 |
| 21 | *TAP1* | 2.02E-18 | 71 | *C5orf56* | 9.10E-08 | 121 | *TAPBP* | 5.62E-06 | 171 | *TCF7* | 3.56E-05 |
| 22 | *SYNE2* | 2.54E-17 | 72 | *NACA* | 9.10E-08 | 122 | *EHD4* | 5.62E-06 | 172 | *GATA3* | 3.56E-05 |
| 23 | *ITGB1* | 3.10E-15 | 73 | *EEF1G* | 1.17E-07 | 123 | *LGALS3* | 5.66E-06 | 173 | *ISG20* | 3.56E-05 |
| 24 | *MX1* | 3.57E-15 | 74 | *TPT1* | 1.32E-07 | 124 | *MX2* | 5.77E-06 | 174 | *REEP5* | 3.66E-05 |
| 25 | *TRIM22* | 5.23E-15 | 75 | *KLF6* | 1.54E-07 | 125 | *ODF2L* | 6.15E-06 | 175 | *GNB1* | 3.84E-05 |

| | | | | | | | | | | | |
|---|---|---|---|---|---|---|---|---|---|---|---|
| 26 | *LY6E* | 6.73E-15 | 76 | *CASP1* | 1.55E-07 | 126 | *PDIA3* | 6.19E-06 | 176 | *CAPN1* | 3.84E-05 |
| 27 | *APOL6* | 1.17E-14 | 77 | *IFIH1* | 1.65E-07 | 127 | *FTL* | 6.24E-06 | 177 | *COTL1* | 3.90E-05 |
| 28 | *S100A11* | 2.67E-14 | 78 | *C12orf57* | 1.71E-07 | 128 | *HLA-E* | 6.64E-06 | 178 | *PHF11* | 3.97E-05 |
| 29 | *NEAT1* | 3.83E-14 | 79 | *CAST* | 2.27E-07 | 129 | *AHR* | 6.72E-06 | 179 | *CALM1* | 4.20E-05 |
| 30 | *XAF1* | 3.89E-14 | 80 | *ANXA2* | 2.35E-07 | 130 | *SQOR* | 6.72E-06 | 180 | *NPM1* | 4.35E-05 |
| 31 | *ISG15* | 6.35E-14 | 81 | *MAF* | 2.51E-07 | 131 | *CD99* | 6.72E-06 | 181 | *GART* | 4.35E-05 |
| 32 | *ANXA1* | 3.05E-12 | 82 | *ADAM19* | 2.81E-07 | 132 | *TXN* | 7.68E-06 | 182 | *EIF4EBP2* | 4.41E-05 |
| 33 | *SH3BP5* | 4.19E-12 | 83 | *MFHAS1* | 2.90E-07 | 133 | *ARPC5L* | 7.70E-06 | 183 | *PFDN5* | 4.53E-05 |
| 34 | *IFI35* | 5.52E-12 | 84 | *FAU* | 2.97E-07 | 134 | *CCDC107* | 9.78E-06 | 184 | *SRGN* | 4.72E-05 |
| 35 | *EIF2AK2* | 6.23E-12 | 85 | *ZBP1* | 3.03E-07 | 135 | *IFITM2* | 1.06E-05 | 185 | *SELENOS* | 4.94E-05 |
| 36 | *EEF1B2* | 1.65E-11 | 86 | *HLA-B* | 3.36E-07 | 136 | *CPQ* | 1.14E-05 | 186 | *OGDH* | 4.96E-05 |
| 37 | *MT2A* | 5.53E-11 | 87 | *SLFN5* | 3.61E-07 | 137 | *ACTR3* | 1.19E-05 | 187 | *PANK2* | 5.00E-05 |
| 38 | *AHNAK* | 6.51E-11 | 88 | *ADAR* | 3.62E-07 | 138 | *LDHA* | 1.22E-05 | 188 | *MRPS27* | 5.21E-05 |
| 39 | *TRIM69* | 1.32E-10 | 89 | *SAMHD1* | 3.72E-07 | 139 | *GSTK1* | 1.23E-05 | 189 | *ALOX5AP* | 5.29E-05 |
| 40 | *TRANK1* | 1.77E-10 | 90 | *MYL12A* | 7.42E-07 | 140 | *PTPN11* | 1.23E-05 | 190 | *EIF4E3* | 5.41E-05 |
| 41 | *TPM3* | 2.64E-10 | 91 | *SAMD9* | 7.47E-07 | 141 | *PFN1* | 1.24E-05 | 191 | *RGS10* | 5.41E-05 |
| 42 | *SH3BGRL3* | 4.16E-10 | 92 | *ACTN4* | 7.56E-07 | 142 | *GPR155* | 1.30E-05 | 192 | *RAB27A* | 5.41E-05 |
| 43 | *PHACTR2* | 5.07E-10 | 93 | *PARP8* | 7.67E-07 | 143 | *LRP10* | 1.33E-05 | 193 | *DDX60* | 5.49E-05 |
| 44 | *HERC5* | 5.16E-10 | 94 | *ZNF267* | 9.10E-07 | 144 | *MYO1G* | 1.38E-05 | 194 | *SRSF5* | 5.49E-05 |
| 45 | *STAT2* | 7.50E-10 | 95 | *HLA-C* | 9.39E-07 | 145 | *CYTOR* | 1.46E-05 | 195 | *YWHAZ* | 5.55E-05 |
| 46 | *IL32* | 7.74E-10 | 96 | *CIRBP* | 1.25E-06 | 146 | *GTF2E2* | 1.48E-05 | 196 | *IRF7* | 5.75E-05 |
| 47 | *APOBEC3G* | 7.74E-10 | 97 | *YWHAH* | 1.27E-06 | 147 | *CCR7* | 1.54E-05 | 197 | *CD7* | 6.01E-05 |
| 48 | *SARAF* | 8.67E-10 | 98 | *S100A10* | 1.30E-06 | 148 | *APOBEC3C* | 1.60E-05 | 198 | *MACF1* | 6.14E-05 |
| 49 | *RNF213* | 1.12E-09 | 99 | *CRIP1* | 1.38E-06 | 149 | *ID2* | 1.62E-05 | 199 | *TLN1* | 6.40E-05 |
| 50 | *S100A4* | 1.16E-09 | 100 | *C4orf48* | 1.39E-06 | 150 | *N4BP1* | 1.71E-05 | 200 | *RNF214* | 6.43E-05 |

**Supplementary Table 11.** Top 20 significant pathways obtained from Enrichr after gene set enrichment of common STAT1 specific SGS-responsive genes from the three PBMC datasets.

| Term | Overlap | P-value | Adj. P-value | Genes |
|---|---|---|---|---|
| *Defense Response To Symbiont (GO:0140546)* | 8/148 | 2.59E-09 | 1.37E-06 | *GBP5;IFI16;OAS2;STAT1;STAT2;NLRC5;EIF2AK2;GBP2* |
| *Defense Response To Virus (GO:0051607)* | 8/189 | 1.78E-08 | 4.68E-06 | *GBP5;IFI16;OAS2;STAT1;STAT2;NLRC5;EIF2AK2;GBP2* |
| *Translational Elongation (GO:0006414)* | 5/42 | 5.69E-08 | 9.99E-06 | *EEF1B2;EEF1A1;EEF1G;EEF1D;RACK1* |
| *Cellular Response To Type II Interferon (GO:0071346)* | 5/66 | 5.72E-07 | 7.53E-05 | *GBP5;SP100;STAT1;GBP2;GBP4* |
| *Response To Type II Interferon (GO:0034341)* | 5/80 | 1.50E-06 | 1.32E-04 | *GBP5;SP100;STAT1;GBP2;GBP4* |
| *Translation (GO:0006412)* | 7/234 | 1.55E-06 | 1.32E-04 | *EEF1B2;EEF1A1;EEF1G;EEF1D;RACK1;FAU;UBA52* |
| *Type I Interferon-Mediated Signaling Pathway (GO:0060337)* | 4/37 | 1.98E-06 | 1.32E-04 | *SP100;STAT1;OAS2;STAT2* |
| *Cellular Response To Type I Interferon (GO:0071357)* | 4/38 | 2.21E-06 | 1.32E-04 | *SP100;STAT1;OAS2;STAT2* |
| *Regulation Of Type II Interferon-Mediated Signaling Pathway (GO:0060334)* | 3/11 | 2.25E-06 | 1.32E-04 | *NLRC5;PARP14;PARP9* |
| *Interferon-Mediated Signaling Pathway (GO:0140888)* | 4/49 | 6.21E-06 | 3.27E-04 | *SP100;OAS2;STAT1;STAT2* |
| *Negative Regulation Of Viral Process (GO:0048525)* | 4/61 | 1.50E-05 | 7.17E-04 | *IFI16;OAS2;STAT1;EIF2AK2* |
| *Response To Interferon-Beta (GO:0035456)* | 3/29 | 4.83E-05 | 2.12E-03 | *IFI16;STAT1;XAF1* |
| *Interleukin-27-Mediated Signaling Pathway (GO:0070106)* | 2/5 | 5.85E-05 | 2.37E-03 | *OAS2;STAT1* |
| *Positive Regulation Of Hydrolase Activity (GO:0051345)* | 5/176 | 6.95E-05 | 2.62E-03 | *DOCK10;PREX1;RACK1;RAPGEF1;SOS1* |
| *Macromolecule Biosynthetic Process (GO:0009059)* | 5/183 | 8.36E-05 | 2.94E-03 | *EEF1B2;EEF1A1;EEF1G;EEF1D;FAU* |
| *Cellular Response To Cytokine Stimulus (GO:0071345)* | 6/308 | 1.02E-04 | 3.35E-03 | *GBP5;IFI16;STAT1;SOS1;GBP2;GBP4* |
| *Response To cAMP (GO:0051591)* | 3/38 | 1.10E-04 | 3.40E-03 | *STAT1;RAPGEF1;AHR* |
| *Positive Regulation Of Cytokine-Mediated Signaling Pathway (GO:0001961)* | 3/44 | 1.71E-04 | 4.83E-03 | *NLRC5;PARP14;PARP9* |
| *Regulation Of GTPase Activity (GO:0043087)* | 5/214 | 1.74E-04 | 4.83E-03 | *DOCK10;PREX1;RACK1;RAPGEF1;SOS1* |
| *Type II Interferon-Mediated Signaling Pathway (GO:0060333)* | 2/9 | 2.09E-04 | 5.52E-03 | *SP100;STAT1* |

**Supplementary Table 12.** Permalinks for STRING interaction analysis using top SGS-responsive genes.

| **Dataset** | **Gene of interest** | **# Top Genes** | **Permalink** | **P-Value** |
|---|---|---|---|---|
| Glioblastoma | *Ccr2* | 50 | https://version-12-0.string-db.org/cgi/network?networkId=bzQTF1kixgVj | 1.00E-16 |
| Spine | *Kdm6b* | 50 | https://version-12-0.string-db.org/cgi/network?networkId=bk0ozYPYuRQ1 | 3.79E-03 |
| PBMC – 20k | *IL7R* | 54 | https://version-12-0.string-db.org/cgi/network?networkId=be3MJw0vAJ4U | 1.17E-12 |
| PBMC | *STAT1* | 49 | https://version-12-0.string-db.org/cgi/network?networkId=bJlDBBzfoUUa | 1.00E-16 |

**Supplementary Table 13.** Gene statistics of target genes from respective datasets using the Spline-HVG method.

| **Dataset** | **Cell type** | **Gene** | **Dropout** | **Log Mean** | **Log CV** | **Distance** | **HVG** |
|---|---|---|---|---|---|---|---|
| Glioblastoma | Monocytes | *Ccr2* | 0.257 | 1.242 | 0.703 | 0.077 | TRUE |
| Embryonic Spine | Motor neurons | *Kdm6b* | 0.477 | 0.715 | 0.885 | 0.131 | TRUE |
| PBMC – 20k | CD4+ T cells | *IL7R* | 0.067 | 2.201 | 0.628 | 0.403 | TRUE |
| PBMC – 5k | CD4+ T cells | *STAT1* | 0.603 | 0.597 | 1.049 | 0.148 | TRUE |
| PBMC – 10k | CD4+ T cells | *STAT1* | 0.458 | 0.831 | 0.899 | 0.198 | TRUE |
| PBMC – 20k | CD4+ T cells | *STAT1* | 0.427 | 0.895 | 0.861 | 0.201 | TRUE |

**Supplementary Table 14.** Comparison of mean total runtimes of scSGS with scTenifoldKnk.

| Number of cells | Number of genes | Mean runtime scTenifoldKnk | Mean runtime scSGS |
|---|---|---|---|
| 500 | 1000 | 48.44 sec | 0.352 sec |
| 1000 | 1000 | 47.80 sec | 0.370 sec |
| 1000 | 3000 | 7 min 29 sec | 0.932 sec |
| 1000 | 5000 | 26 min 18 sec | 1.570 sec |
| 3000 | 5000 | 26 min 28 sec | 2.012 sec |

#### Supplementary Figures

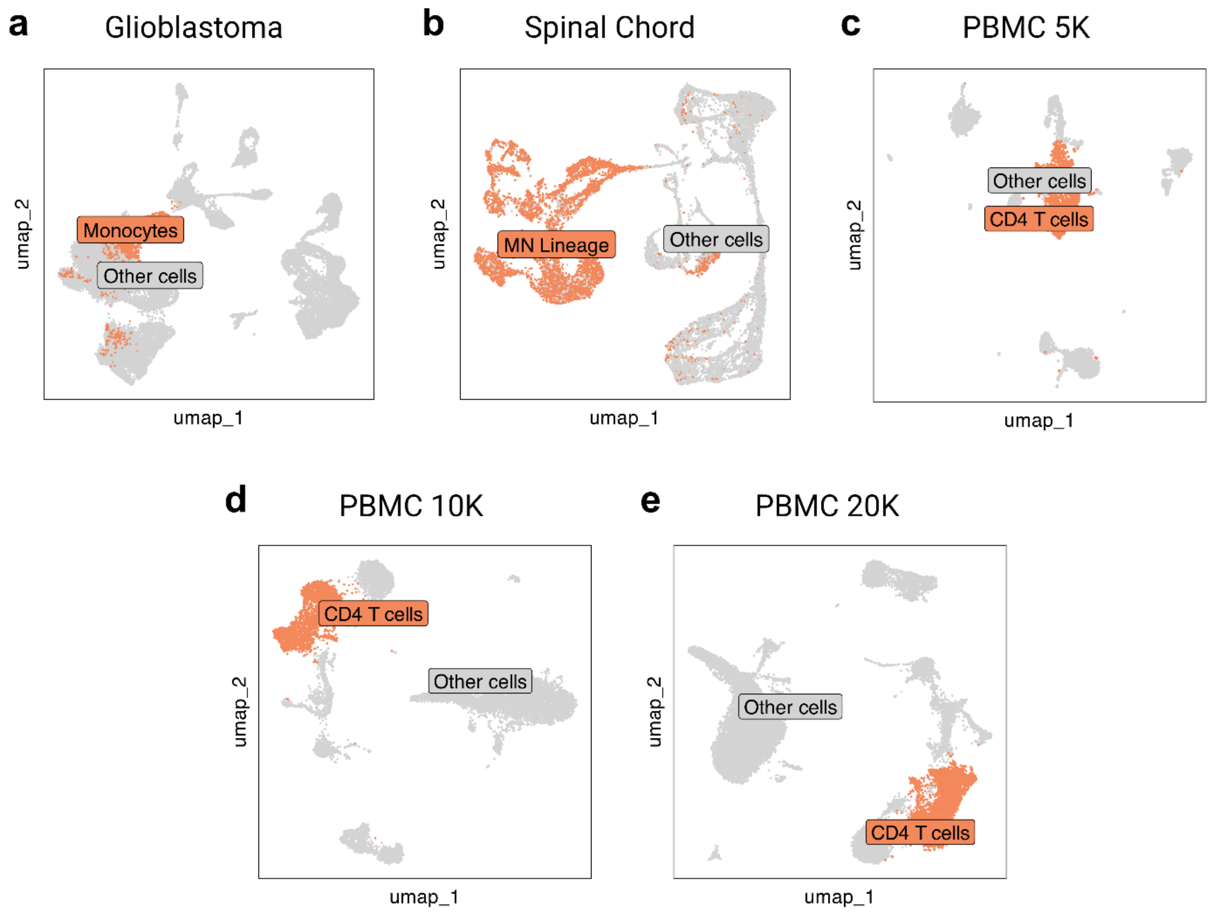

**Supplementary Fig. 1.** UMAP plots highlighting cell types/states used for scSGS analysis from the respective datasets.

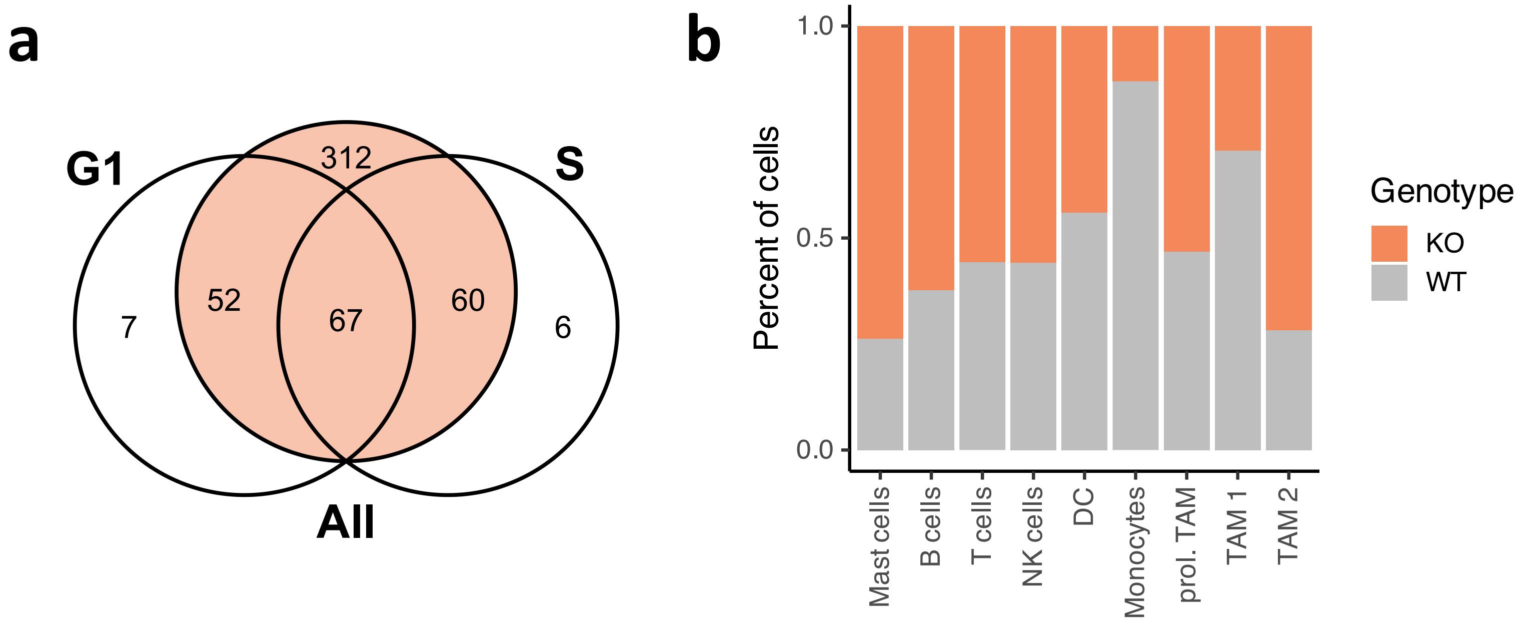

**Supplementary Fig. 2.** a) Overlap of SGS-responsive genes identified from all WT monocytes, G1 phase monocytes, and S phase monocytes. b) Cell percentages across cell types in WT and *Ccr2* KO samples.

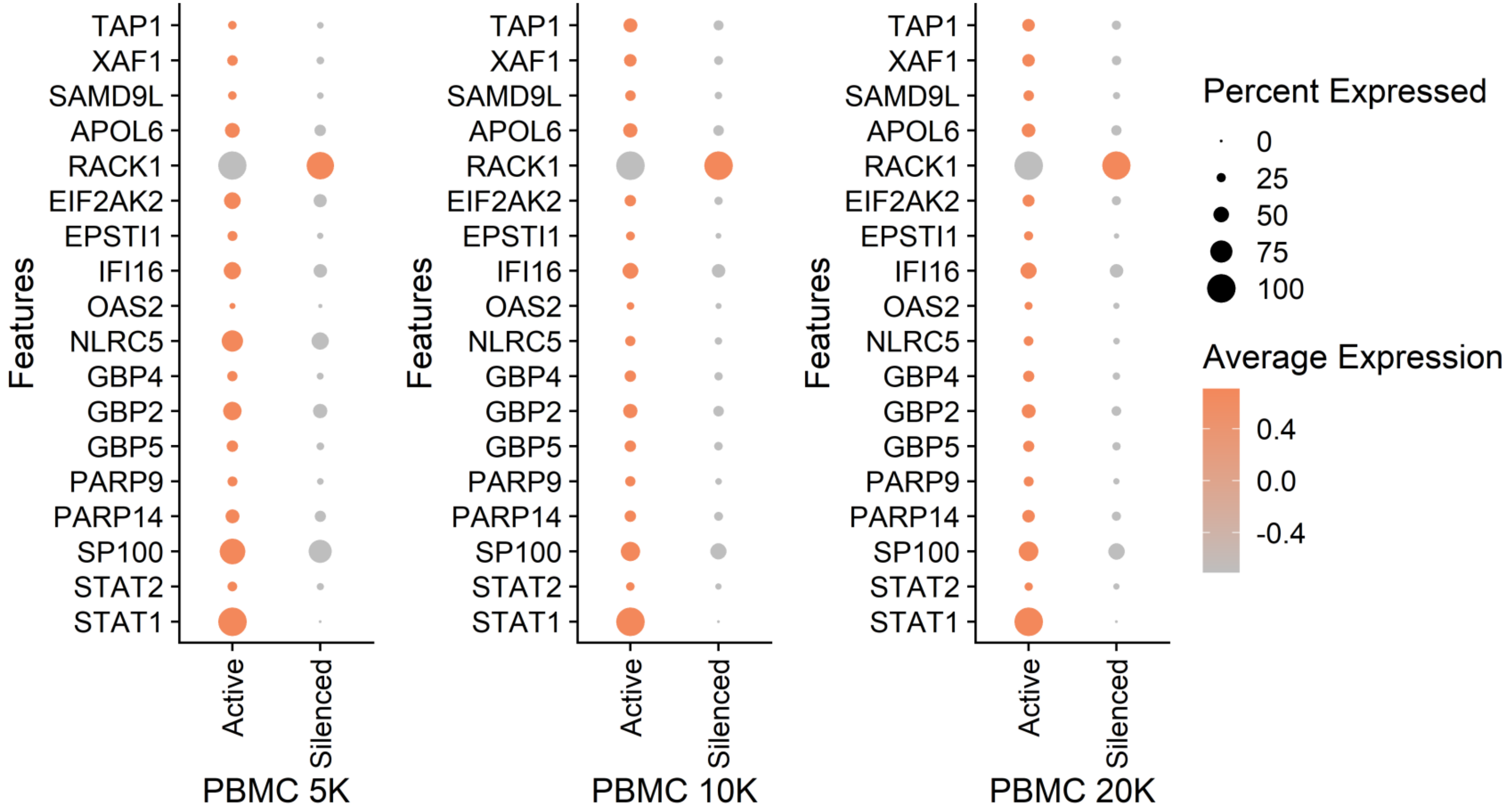

**Supplementary Fig. 3.** Expressions of SGS-responsive genes directly linked with STAT1 according to STRING interaction analysis in the three PBMC datasets in Active and Silenced subsets.

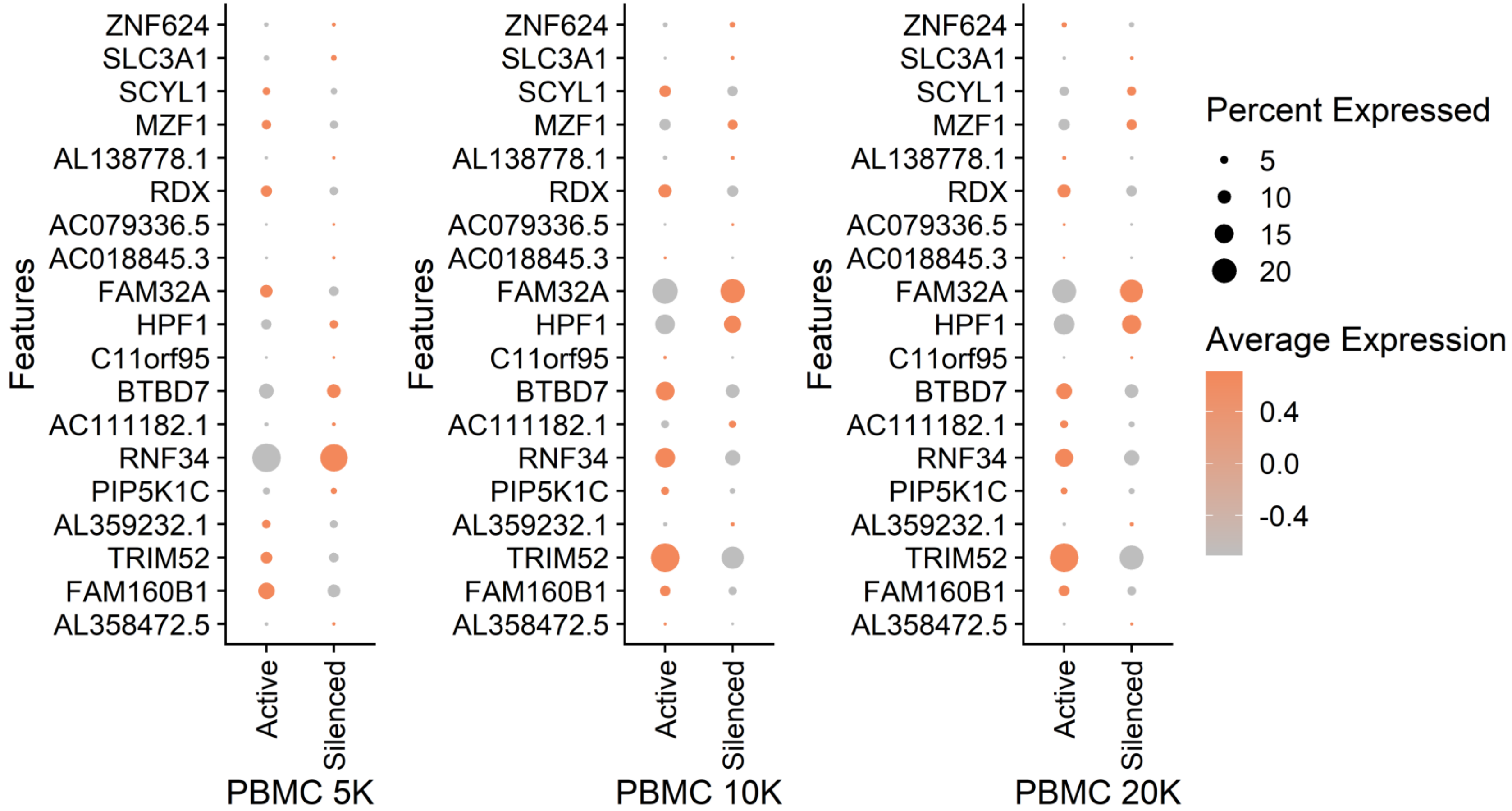

**Supplementary Fig. 4.** Expressions of 18 randomly selected genes from the three PBMC datasets in Active and Silenced subsets.

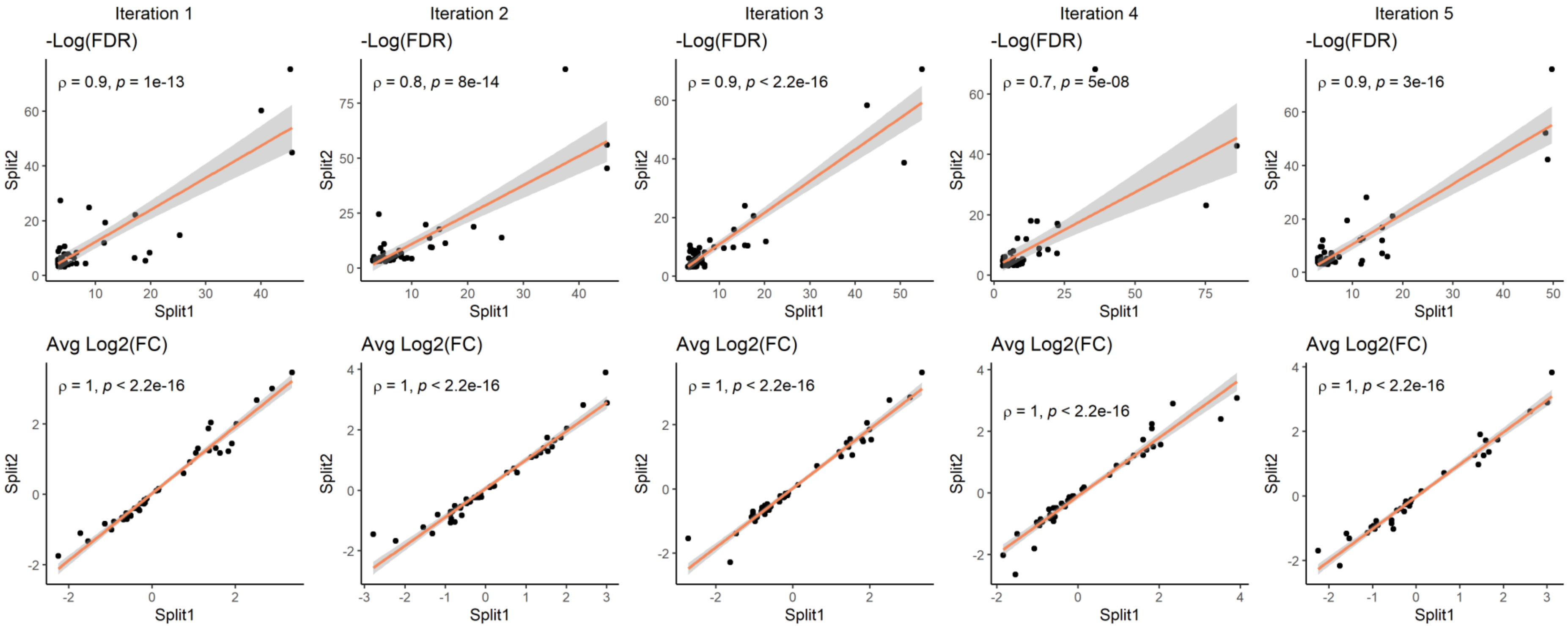

**Supplementary Fig. 5.** Relationship between FDR and average Log2(FC) values between two randomly split samples from the same PBM20K dataset over 5 iterations.

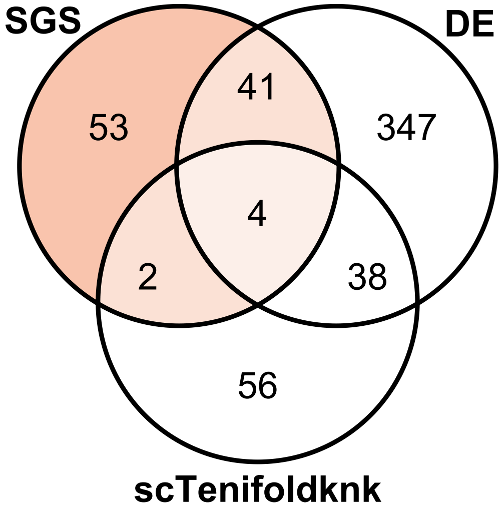

**Supplementary Fig. 6.** Overlap of top 100 SGS-responsive and top 100 significant virtual KO genes from scTenifoldKnk with top 100 DE genes from *in vivo Ccr2* KO study

5 20k Human PBMCs, 3' HT v3.1, Chromium X, Single Cell Gene Expression Dataset by Cell Ranger 6.1.0, 10x Genomics. (2021, August 9).
